## Supplemental Material for "Genomic basis of drought resistance in *Fagus sylvatica*"

**Suppl. Table 1**. Sampled *Fagus sylvatica* individuals. Given are the phenotype, sampling location, sampling date, geographical coordinates in decimal degrees, the pool to which the individual contributed to (h=healthy, d=damaged), whether it was individually re-sequenced (x) and whether it was used in the SNP assay to validate the results (x).

| TreeID | Phenotype | Location | Sampling date | Latitude | Longitude | Pool (set 1) | individually re-sequenced | Validation (set 2) |
|---|---|---|---|---|---|---|---|---|
| PF_001 | damaged | Königstein | 29.08.2019 | 50.1957342 | 8.4648591 | dNorth | | |
| PF_002 | healthy | Königstein | 29.08.2019 | 50.1957187 | 8.4648341 | hNorth | x | |
| PF_003 | damaged | Königstein | 29.08.2019 | 50.1964726 | 8.4664262 | dNorth | x | |
| PF_004 | healthy | Königstein | 29.08.2019 | 50.1964232 | 8.4664275 | hNorth | x | |
| PF_005 | damaged | Königstein | 29.08.2019 | 50.1929421 | 8.456941 | dNorth | x | |
| PF_006 | healthy | Königstein | 29.08.2019 | 50.1928331 | 8.4568086 | hNorth | x | |
| PF_007 | damaged | Königstein | 29.08.2019 | 50.1925453 | 8.457816 | dNorth | x | |
| PF_008 | healthy | Königstein | 29.08.2019 | 50.1925741 | 8.4578839 | hNorth | x | |
| PF_009 | damaged | Königstein | 29.08.2019 | 50.1926029 | 8.4579518 | dNorth | x | |
| PF_010 | healthy | Königstein | 29.08.2019 | 50.1926317 | 8.4580197 | hNorth | x | |
| PF_011 | damaged | Obernhain | 03.09.2019 | 50.2825898 | 8.5712681 | dNorth | | |
| PF_012 | healthy | Obernhain | 03.09.2019 | 50.2827911 | 8.5711075 | hNorth | | |
| PF_013 | damaged | Obernhain | 03.09.2019 | 50.2820637 | 8.5734056 | dNorth | | |
| PF_014 | healthy | Obernhain | 03.09.2019 | 50.282214 | 8.5734601 | hNorth | | |
| PF_015 | damaged | Obernhain | 03.09.2019 | 50.2817116 | 8.5761923 | dNorth | | |
| PF_016 | healthy | Obernhain | 03.09.2019 | 50.2816381 | 8.5761487 | hNorth | x | |
| PF_017 | damaged | Obernhain | 03.09.2019 | 50.2821782 | 8.5704083 | dNorth | x | |
| PF_018 | healthy | Obernhain | 03.09.2019 | 50.2821327 | 8.5704883 | hNorth | x | |
| PF_019 | damaged | Obernhain | 03.09.2019 | 50.2816941 | 8.5700475 | dNorth | x | |
| PF_020 | healthy | Obernhain | 03.09.2019 | 50.2817058 | 8.570133 | hNorth | x | |
| PF_021 | damaged | Usingen | 04.09.2019 | 50.3392732 | 8.4913495 | dNorth | x | |
| PF_022 | healthy | Usingen | 04.09.2019 | 50.3391762 | 8.491312 | hNorth | x | |
| PF_023 | damaged | Usingen | 04.09.2019 | 50.3397337 | 8.4924001 | dNorth | x | |
| PF_024 | healthy | Usingen | 04.09.2019 | 50.3396942 | 8.4924593 | hNorth | x | |
| PF_025 | damaged | Usingen | 04.09.2019 | 50.344079 | 8.4932305 | dNorth | x | |
| PF_026 | healthy | Usingen | 04.09.2019 | 50.3440912 | 8.4933098 | hNorth | x | |
| PF_027 | damaged | Usingen | 04.09.2019 | 50.3452681 | 8.4959808 | dNorth | x | |
| PF_028 | healthy | Usingen | 04.09.2019 | 50.3451246 | 8.4962431 | hNorth | x | |
| PF_029 | damaged | Usingen | 04.09.2019 | 50.3416761 | 8.4918323 | dNorth | x | |
| PF_030 | healthy | Usingen | 04.09.2019 | 50.3417203 | 8.4917173 | hNorth | x | |
| PF_031 | damaged | Eppenhain | 05.09.2019 | 50.1736813 | 8.3908577 | dNorth | x | |

| | | | | | | | |
|---|---|---|---|---|---|---|---|
| PF_032 | healthy | Eppenhain | 05.09.2019 | 50.1736578 | 8.39113553 | hNorth | x |
| PF_033 | damaged | Eppenhain | 05.09.2019 | 50.1750866 | 8.3891472 | dNorth | x |
| PF_034 | healthy | Eppenhain | 05.09.2019 | 50.175069 | 8.3892187 | hNorth | x |
| PF_035 | damaged | Eppenhain | 05.09.2019 | 50.1754443 | 8.3876407 | dNorth | x |
| PF_036 | healthy | Eppenhain | 05.09.2019 | 50.1754007 | 8.3875849 | hNorth | x |
| PF_037 | damaged | Eppenhain | 05.09.2019 | 50.1746449 | 8.3872509 | dNorth | x |
| PF_038 | healthy | Eppenhain | 05.09.2019 | 50.1746095 | 8.3874874 | hNorth | x |
| PF_039 | damaged | Eppenhain | 05.09.2019 | 50.1734334 | 8.3904983 | dNorth | x |
| PF_040 | healthy | Eppenhain | 05.09.2019 | 50.1733173 | 8.390211 | hNorth | x |
| PF_041 | damaged | Eppenhain | 05.09.2019 | 50.1745915 | 8.3907433 | dNorth | x |
| PF_042 | healthy | Eppenhain | 05.09.2019 | 50.1745128 | 8.3908124 | hNorth | x |
| PF_050 | damaged | Bad Camberg | 06.09.2019 | 50.2976336 | 8.2980473 | dNorth | |
| PF_051 | healthy | Bad Camberg | 06.09.2019 | 50.2975811 | 8.298169 | hNorth | |
| PF_052 | damaged | Daubringen | 07.09.2019 | 50.6483759 | 8.7419719 | dNorth | x |
| PF_053 | healthy | Daubringen | 07.09.2019 | 50.6482664 | 8.7421457 | hNorth | x |
| PF_054 | damaged | Daubringen | 07.09.2019 | 50.6480228 | 8.7428739 | dNorth | x |
| PF_055 | healthy | Daubringen | 07.09.2019 | 50.6478157 | 8.742954 | hNorth | x |
| PF_056 | damaged | Daubringen | 07.09.2019 | 50.648846 | 8.7426283 | dNorth | x |
| PF_057 | healthy | Daubringen | 07.09.2019 | 50.6489798 | 8.7423211 | hNorth | x |
| PF_058 | damaged | Schwabendorf | 07.09.2019 | 50.8927904 | 8.8879419 | dNorth | x |
| PF_059 | healthy | Schwabendorf | 07.09.2019 | 50.8929008 | 8.8881284 | hNorth | x |
| PF_060 | damaged | Schwabendorf | 07.09.2019 | 50.8951542 | 8.8822243 | dNorth | x |
| PF_061 | healthy | Schwabendorf | 07.09.2019 | 50.8952005 | 8.882144 | hNorth | x |
| PF_062 | damaged | Oberurff | 07.09.2019 | 51.0458016 | 9.1505195 | dNorth | x |
| PF_063 | healthy | Oberurff | 07.09.2019 | 51.0457499 | 9.1505234 | hNorth | x |
| PF_064 | damaged | Oberurff | 07.09.2019 | 51.0461483 | 9.1478112 | dNorth | x |
| PF_065 | healthy | Oberurff | 07.09.2019 | 51.0461706 | 9.1474544 | hNorth | x |
| PF_066 | damaged | Oberurff | 07.09.2019 | 51.0453396 | 9.1466057 | dNorth | x |
| PF_067 | healthy | Oberurff | 07.09.2019 | 51.0453396 | 9.146057 | hNorth | x |
| PF_068 | damaged | Oberurff | 07.09.2019 | 51.0463978 | 9.1432389 | dNorth | x |
| PF_069 | healthy | Oberurff | 07.09.2019 | 51.0464672 | 9.1435546 | hNorth | x |
| PF_070 | damaged | Neukirchen | 08.09.2019 | 50.8848091 | 9.3557289 | dNorth | x |
| PF_071 | healthy | Neukirchen | 08.09.2019 | 50.88496 | 9.3560291 | hNorth | x |
| PF_072 | damaged | Langenhain | 09.09.2019 | 50.0955818 | 8.4101217 | dNorth | x |
| PF_073 | healthy | Langenhain | 09.09.2019 | 50.0956666 | 8.4105235 | hNorth | x |
| PF_074 | damaged | Langenhain | 09.09.2019 | 50.0959896 | 8.4104922 | dNorth | x |
| PF_075 | healthy | Langenhain | 09.09.2019 | 50.0960509 | 8.4105981 | hNorth | x |
| PF_076 | damaged | Langenhain | 09.09.2019 | 50.0947953 | 8.4151317 | dNorth | x |

| | | | | | | | |
|---|---|---|---|---|---|---|---|
| PF_077 | healthy | Langenhain | 09.09.2019 | 50.0947434 | 8.4150994 | hNorth | x |
| PF_078 | damaged | Langenhain | 09.09.2019 | 50.0911467 | 8.41578726 | dNorth | x |
| PF_079 | healthy | Langenhain | 09.09.2019 | 50.0912627 | 8.415817 | hNorth | |
| PF_080 | damaged | Langenhain | 09.09.2019 | 50.0901299 | 8.4166724 | dNorth | x |
| PF_081 | healthy | Langenhain | 09.09.2019 | 50.090096 | 8.4166748 | hNorth | x |
| PF_082 | damaged | Langenhain | 09.09.2019 | 50.0940815 | 8.4100466 | dNorth | x |
| PF_083 | healthy | Langenhain | 09.09.2019 | 50.0939183 | 8.4099548 | hNorth | x |
| PF_085 | damaged | Theistal | 11.09.2019 | 50.1480865 | 8.2445088 | dNorth | x |
| PF_086 | healthy | Theistal | 11.09.2019 | 50.1480164 | 8.2447881 | hNorth | x |
| PF_087 | damaged | Theistal | 11.09.2019 | 50.1517267 | 8.2464193 | dNorth | x |
| PF_088 | healthy | Theistal | 11.09.2019 | 50.151875 | 8.2461256 | hNorth | x |
| PF_089 | damaged | Theistal | 11.09.2019 | 50.1532534 | 8.2456221 | dNorth | x |
| PF_090 | healthy | Theistal | 11.09.2019 | 50.1528409 | 8.2455616 | hNorth | x |
| PF_091 | damaged | Theistal | 11.09.2019 | 50.1539513 | 8.2466804 | dNorth | x |
| PF_092 | healthy | Theistal | 11.09.2019 | 50.1540134 | 8.2462949 | hNorth | x |
| PF_093 | damaged | Schlangenbad | 11.09.2019 | 50.1027276 | 8.1136673 | dNorth | x |
| PF_094 | healthy | Schlangenbad | 11.09.2019 | 50.1025753 | 8.1136931 | hNorth | x |
| PF_095 | damaged | Schlangenbad | 11.09.2019 | 50.0997272 | 8.1101568 | dNorth | x |
| PF_096 | healthy | Schlangenbad | 11.09.2019 | 50.099823 | 8.1103387 | hNorth | x |
| PF_100 | damaged | Wehrheim | 12.09.2019 | 50.3153263 | 8.5819508 | dNorth | x |
| PF_101 | healthy | Wehrheim | 12.09.2019 | 50.315372 | 8.5820711 | hNorth | x |
| PF_102 | damaged | Wehrheim | 12.09.2019 | 50.3145513 | 8.5854558 | dNorth | x |
| PF_103 | healthy | Wehrheim | 12.09.2019 | 50.3145748 | 8.585572 | hNorth | x |
| PF_104 | damaged | Wehrheim | 12.09.2019 | 50.3147404 | 8.585804 | dNorth | x |
| PF_105 | healthy | Wehrheim | 12.09.2019 | 50.3146745 | 8.5858773 | hNorth | x |
| PF_106 | damaged | Wehrheim | 12.09.2019 | 50.3149149 | 8.5859893 | dNorth | x |
| PF_107 | healthy | Wehrheim | 12.09.2019 | 50.31472242 | 8.5861579 | hNorth | x |
| PF_108 | damaged | Wehrheim | 12.09.2019 | 50.3149559 | 8.5866045 | dNorth | x |
| PF_109 | healthy | Wehrheim | 12.09.2019 | 50.3150733 | 8.5866527 | hNorth | x |
| PF_110 | damaged | Wehrheim | 12.09.2019 | 50.3155392 | 8.587186 | dNorth | x |
| PF_111 | healthy | Wehrheim | 12.09.2019 | 50.3154613 | 8.5872255 | hNorth | x |
| PF_112 | damaged | Wehrheim | 12.09.2019 | 50.3130127 | 8.580265 | dNorth | x |
| PF_113 | healthy | Wehrheim | 12.09.2019 | 50.3130103 | 8.5800785 | hNorth | x |
| PF_114 | damaged | Wehrheim | 12.09.2019 | 50.3141899 | 8.581059 | dNorth | x |
| PF_115 | healthy | Wehrheim | 12.09.2019 | 50.314164 | 8.5809636 | hNorth | x |
| S_001 | damaged | Bot. Garten | 26.08.2019 | 49.79971 | 8.95682 | dSouth | |
| S_002 | healthy | Bot. Garten | 26.08.2019 | 49.86852 | 8.67999 | hSouth | |
| S_003 | damaged | Bot. Garten | 26.08.2019 | 49.86856 | 8.68018 | dSouth | |

| | | | | | | |
|---|---|---|---|---|---|---|
| S_004 | healthy | Bot. Garten | 26.08.2019 | 49.86856 | 8.68018 | hSouth |
| S_005 | damaged | Vivarium | 26.08.2019 | 49.86855 | 8.68017 | dSouth |
| S_006 | healthy | Vivarium | 26.08.2019 | 49.86436 | 8.6842 | hSouth |
| S_007 | damaged | Vivarium | 26.08.2019 | 49.8644 | 8.68418 | dSouth |
| S_008 | healthy | Vivarium | 26.08.2019 | 49.8644 | 8.68418 | hSouth |
| S_009 | damaged | Vivarium | 26.08.2019 | 49.8644 | 8.68418 | dSouth |
| S_010 | healthy | Vivarium | 26.08.2019 | 49.8644 | 8.68418 | hSouth |
| S_011 | damaged | Vivarium | 26.08.2019 | 49.8644 | 8.68418 | dSouth |
| S_012 | healthy | Vivarium | 26.08.2019 | 49.8644 | 8.68418 | hSouth |
| S_013 | damaged | VivariumII | 26.08.2019 | 49.8644 | 8.68418 | dSouth |
| S_014 | healthy | VivariumII | 26.08.2019 | 49.8644 | 8.68418 | hSouth |
| S_015 | damaged | VivariumII | 26.08.2019 | 49.86227 | 8.68876 | dSouth |
| S_016 | healthy | VivariumII | 26.08.2019 | 49.86241 | 8.68874 | hSouth |
| S_017 | damaged | VivariumII | 26.08.2019 | 49.86199 | 8.68894 | dSouth |
| S_018 | healthy | VivariumII | 26.08.2019 | 49.86196 | 8.68885 | hSouth |
| S_019 | damaged | VivariumII | 26.08.2019 | 49.86092 | 8.69278 | dSouth |
| S_020 | healthy | VivariumII | 26.08.2019 | 49.86031 | 8.69225 | hSouth |
| S_021 | damaged | VivariumIII | 27.08.2019 | 49.85883 | 8.69445 | dSouth |
| S_022 | healthy | VivariumIII | 27.08.2019 | 49.85883 | 8.69445 | hSouth |
| S_023 | damaged | VivariumIII | 27.08.2019 | 49.85882 | 8.69446 | dSouth |
| S_024 | healthy | VivariumIII | 27.08.2019 | 49.85882 | 8.69446 | hSouth |
| S_025 | damaged | VivariumIII | 27.08.2019 | 49.85862 | 8.6944 | dSouth |
| S_026 | healthy | VivariumIII | 27.08.2019 | 49.85862 | 8.6944 | hSouth |
| S_027 | damaged | VivariumIII | 27.08.2019 | 49.85637 | 8.69584 | dSouth |
| S_028 | healthy | VivariumIII | 27.08.2019 | 49.85637 | 8.69584 | hSouth |
| S_029 | damaged | VivariumIII | 27.08.2019 | 49.8565 | 8.69616 | dSouth |
| S_030 | healthy | VivariumIII | 27.08.2019 | 49.8565 | 8.69616 | hSouth |
| S_031 | damaged | VivariumIII | 27.08.2019 | 49.8565 | 8.69616 | dSouth |
| S_032 | healthy | VivariumIII | 27.08.2019 | 49.8565 | 8.69616 | hSouth |
| S_033 | damaged | VivariumIII | 27.08.2019 | 49.85576 | 8.69886 | dSouth |
| S_034 | healthy | VivariumIII | 27.08.2019 | 49.85576 | 8.69886 | hSouth |
| S_035 | damaged | VivariumIII | 27.08.2019 | 49.85576 | 8.69886 | dSouth |
| S_036 | healthy | VivariumIII | 27.08.2019 | 49.85576 | 8.69886 | hSouth |
| S_037 | damaged | VivariumIII | 27.08.2019 | 49.85329 | 8.6938 | dSouth |
| S_038 | healthy | VivariumIII | 27.08.2019 | 49.85329 | 8.6938 | hSouth |
| S_039 | damaged | VivariumIII | 27.08.2019 | 49.85329 | 8.6938 | dSouth |
| S_040 | healthy | VivariumIII | 27.08.2019 | 49.85329 | 8.6938 | hSouth |
| S_041 | damaged | VivariumIII | 27.08.2019 | 49.85021 | 8.9391 | dSouth |

| | | | | | | | |
|---|---|---|---|---|---|---|---|
| S_042 | healthy | VivariumIII | 27.08.2019 | 49.85021 | 8.9391 | hSouth | |
| S_043 | damaged | Neu-Isenburg | 28.08.2019 | 49.85585 | 8.69554 | dSouth | |
| S_044 | healthy | Neu-Isenburg | 28.08.2019 | 49.85585 | 8.69554 | hSouth | |
| S_045 | damaged | Neu-Isenburg | 28.08.2019 | 49.85585 | 8.69554 | dSouth | |
| S_046 | healthy | Neu-Isenburg | 28.08.2019 | 49.85585 | 8.69554 | hSouth | |
| S_047 | damaged | Neu-Isenburg | 28.08.2019 | 49.85585 | 8.69554 | dSouth | |
| S_048 | healthy | Neu-Isenburg | 28.08.2019 | 49.85585 | 8.69554 | hSouth | |
| S_049 | damaged | Neu-Isenburg | 28.08.2019 | 49.85585 | 8.69554 | dSouth | |
| S_050 | healthy | Neu-Isenburg | 28.08.2019 | 49.85585 | 8.69554 | hSouth | |
| S_051 | damaged | Neu-Isenburg | 28.08.2019 | 49.85585 | 8.69554 | dSouth | |
| S_052 | healthy | Neu-Isenburg | 28.08.2019 | 49.85585 | 8.69554 | hSouth | |
| S_053 | damaged | Neu-Isenburg | 28.08.2019 | 49.85585 | 8.69554 | dSouth | |
| S_054 | healthy | Neu-Isenburg | 28.08.2019 | 49.85585 | 8.69554 | hSouth | |
| S_055 | damaged | Neu-Isenburg | 28.08.2019 | 49.85585 | 8.69554 | dSouth | |
| S_056 | healthy | Neu-Isenburg | 28.08.2019 | 49.85585 | 8.69554 | hSouth | |
| S_057 | damaged | Neu-Isenburg | 28.08.2019 | 49.85585 | 8.69554 | dSouth | |
| S_058 | healthy | Neu-Isenburg | 28.08.2019 | 49.85585 | 8.69554 | hSouth | |
| S_059 | damaged | Neu-Isenburg | 28.08.2019 | 49.85585 | 8.69554 | dSouth | |
| S_060 | healthy | Neu-Isenburg | 28.08.2019 | 49.85585 | 8.69554 | hSouth | |
| S_061 | damaged | Westwald | 29.09.2019 | 49.855069 | 8.617017 | dSouth | |
| S_062 | healthy | Westwald | 29.09.2019 | 49.855069 | 8.617017 | hSouth | |
| S_063 | damaged | Westwald | 29.09.2019 | 49.85495 | 8.61703 | dSouth | x |
| S_064 | healthy | Westwald | 29.09.2019 | 49.85495 | 8.61703 | hSouth | |
| S_065 | damaged | Westwald | 29.09.2019 | 49.85495 | 8.61703 | dSouth | |
| S_066 | healthy | Westwald | 29.09.2019 | 49.85495 | 8.61703 | hSouth | x |
| S_067 | damaged | Westwald | 29.09.2019 | 49.85494 | 8.61702 | dSouth | |
| S_068 | healthy | Westwald | 29.09.2019 | 49.85494 | 8.61702 | hSouth | |
| S_069 | damaged | Westwald | 29.09.2019 | 49.854909 | 8.618146 | dSouth | |
| S_070 | healthy | Westwald | 29.09.2019 | 49.854909 | 8.618146 | hSouth | |
| S_071 | damaged | Westwald | 29.09.2019 | 49.85494 | 8.61702 | dSouth | x |
| S_072 | healthy | Westwald | 29.09.2019 | 49.85494 | 8.61702 | hSouth | x |
| S_073 | damaged | Westwald | 29.09.2019 | 49.85494 | 8.61702 | dSouth | |
| S_074 | healthy | Westwald | 29.09.2019 | 49.85494 | 8.61702 | hSouth | |
| S_075 | damaged | Westwald | 29.09.2019 | 49.85494 | 8.61702 | dSouth | |
| S_076 | healthy | Westwald | 29.09.2019 | 49.85494 | 8.61702 | hSouth | |
| S_077 | damaged | Westwald | 29.09.2019 | 49.85494 | 8.61702 | dSouth | x |
| S_078 | healthy | Westwald | 29.09.2019 | 49.85494 | 8.61702 | hSouth | x |
| S_079 | damaged | Westwald | 29.09.2019 | 49.85494 | 8.61702 | dSouth | |

| | | | | | | | | |
|---|---|---|---|---|---|---|---|---|
| S_080 | healthy | Westwald | 29.09.2019 | 49.85494 | 8.61702 | hSouth | | |
| S_081 | damaged | Westwald | 29.09.2019 | 49.85494 | 8.61702 | dSouth | | X |
| S_082 | healthy | Westwald | 29.09.2019 | 49.85494 | 8.61702 | hSouth | | X |
| S_083 | damaged | Westwald | 29.09.2019 | 49.85494 | 8.61702 | dSouth | | |
| S_084 | healthy | Westwald | 29.09.2019 | 49.85494 | 8.61702 | hSouth | | |
| S_085 | damaged | Roßdorf | 02.09.2019 | 49.85494 | 8.61702 | dSouth | | |
| S_086 | healthy | Roßdorf | 02.09.2019 | 49.85494 | 8.61702 | hSouth | | |
| S_087 | damaged | Roßdorf | 02.09.2019 | 49.85943 | 8.70791 | dSouth | | X |
| S_088 | healthy | Roßdorf | 02.09.2019 | 49.85943 | 8.70791 | hSouth | | X |
| S_089 | damaged | Roßdorf | 02.09.2019 | 49.85943 | 8.70786 | dSouth | | |
| S_090 | healthy | Roßdorf | 02.09.2019 | 49.85943 | 8.70786 | hSouth | | |
| S_091 | damaged | Roßdorf | 02.09.2019 | 49.85977 | 8.70795 | dSouth | | |
| S_092 | healthy | Roßdorf | 02.09.2019 | 49.85977 | 8.70795 | hSouth | | |
| S_093 | damaged | Roßdorf | 02.09.2019 | 49.86221 | 8.718 | dSouth | | X |
| S_094 | healthy | Roßdorf | 02.09.2019 | 49.86221 | 8.718 | hSouth | | X |
| S_095 | damaged | Roßdorf | 02.09.2019 | 49.86211 | 8.71801 | dSouth | | |
| S_096 | healthy | Roßdorf | 02.09.2019 | 49.86211 | 8.71801 | hSouth | | |
| S_097 | damaged | Roßdorf | 02.09.2019 | 49.86211 | 8.71801 | dSouth | | |
| S_098 | healthy | Roßdorf | 02.09.2019 | 49.86211 | 8.71801 | hSouth | | |
| S_099 | damaged | Roßdorf | 02.09.2019 | 49.86089 | 8.70835 | dSouth | | |
| S_100 | healthy | Roßdorf | 02.09.2019 | 49.86089 | 8.70835 | hSouth | | |
| S_101 | damaged | Roßdorf | 02.09.2019 | 49.86095 | 8.70771 | dSouth | | |
| S_102 | healthy | Roßdorf | 02.09.2019 | 49.86095 | 8.70771 | hSouth | | |
| S_103 | damaged | Roßdorf | 02.09.2019 | 49.86103 | 8.70769 | | | x |
| S_104 | healthy | Roßdorf | 02.09.2019 | 49.86103 | 8.70769 | | | x |
| S_105 | damaged | Burg Breuberg | 03.09.2019 | 49.8259 | 9.0341 | dSouth | | |
| S_106 | healthy | Burg Breuberg | 03.09.2019 | 49.8259 | 9.0341 | hSouth | | |
| S_107 | damaged | Burg Breuberg | 03.09.2019 | 49.82447 | 9.03196 | dSouth | | |
| S_108 | healthy | Burg Breuberg | 03.09.2019 | 49.82447 | 9.03196 | hSouth | | |
| S_109 | damaged | Burg Breuberg | 03.09.2019 | 49.82457 | 9.03155 | dSouth | | |
| S_110 | healthy | Burg Breuberg | 03.09.2019 | 49.82457 | 9.03155 | hSouth | | |
| S_111 | damaged | Burg Breuberg | 03.09.2019 | 49.82736 | 9.0267 | dSouth | | |
| S_112 | healthy | Burg Breuberg | 03.09.2019 | 49.82736 | 9.0267 | hSouth | | |
| S_113 | damaged | Burg Breuberg | 03.09.2019 | 49.827 | 9.02706 | dSouth | | |
| S_114 | healthy | Burg Breuberg | 03.09.2019 | 49.827 | 9.02706 | hSouth | | |
| S_115 | damaged | Burg Breuberg | 03.09.2019 | 49.82689 | 9.02076 | dSouth | | |
| S_116 | healthy | Burg Breuberg | 03.09.2019 | 49.82689 | 9.02076 | hSouth | | |
| S_117 | damaged | Burg Breuberg | 03.09.2019 | 49.82692 | 9.02038 | | | x |

| | | | | | | | |
|---|---|---|---|---|---|---|---|
| S_118 | healthy | Burg Breuberg | 03.09.2019 | 49.82692 | 9.02038 | | x |
| S_119 | damaged | Burg Breuberg | 03.09.2019 | 49.82322 | 9.02107 | dSouth | |
| S_120 | healthy | Burg Breuberg | 03.09.2019 | 49.82322 | 9.02107 | hSouth | |
| S_121 | damaged | Burg Breuberg | 03.09.2019 | 49.82305 | 9.02048 | dSouth | |
| S_122 | healthy | Burg Breuberg | 03.09.2019 | 49.82305 | 9.02048 | hSouth | |
| S_123 | damaged | Burg Breuberg | 03.09.2019 | 49.82301 | 9.02096 | dSouth | |
| S_124 | healthy | Burg Breuberg | 03.09.2019 | 49.82301 | 9.02096 | hSouth | |
| S_125 | damaged | Heusenstamm | 04.09.2019 | 50.04919 | 8.69445 | dSouth | |
| S_126 | healthy | Heusenstamm | 04.09.2019 | 50.04919 | 8.69445 | hSouth | |
| S_127 | damaged | Heusenstamm | 04.09.2019 | 50.04919 | 8.69445 | dSouth | |
| S_128 | healthy | Heusenstamm | 04.09.2019 | 50.04919 | 8.69445 | hSouth | |
| S_129 | damaged | Heusenstamm | 04.09.2019 | 50.0577 | 8.7788 | | x |
| S_130 | healthy | Heusenstamm | 04.09.2019 | 50.0577 | 8.7788 | | x |
| S_131 | damaged | Heusenstamm | 04.09.2019 | 50.0575 | 8.7787 | dSouth | |
| S_132 | healthy | Heusenstamm | 04.09.2019 | 50.0575 | 8.7787 | hSouth | |
| S_133 | damaged | Heusenstamm | 04.09.2019 | 50.05774 | 8.7796 | dSouth | |
| S_134 | healthy | Heusenstamm | 04.09.2019 | 50.05774 | 8.7796 | hSouth | |
| S_135 | damaged | Heusenstamm | 04.09.2019 | 50.05574 | 8.7795 | dSouth | |
| S_136 | healthy | Heusenstamm | 04.09.2019 | 50.05574 | 8.7795 | hSouth | |
| S_137 | damaged | Heusenstamm | 04.09.2019 | 50.0562 | 8.7787 | dSouth | |
| S_138 | healthy | Heusenstamm | 04.09.2019 | 50.0562 | 8.7787 | hSouth | |
| S_139 | damaged | Heusenstamm | 04.09.2019 | 50.0562 | 8.7787 | dSouth | |
| S_140 | healthy | Heusenstamm | 04.09.2019 | 50.0562 | 8.7787 | hSouth | |
| S_141 | damaged | Heusenstamm | 04.09.2019 | 50.0562 | 8.7787 | dSouth | |
| S_142 | healthy | Heusenstamm | 04.09.2019 | 50.0562 | 8.7787 | hSouth | |
| S_143 | damaged | Heusenstamm | 04.09.2019 | 50.0562 | 8.7787 | | x |
| S_144 | healthy | Heusenstamm | 04.09.2019 | 50.0562 | 8.7787 | | x |
| S_145 | damaged | Heusenstamm | 04.09.2019 | 50.0556 | 8.7788 | | x |
| S_146 | healthy | Heusenstamm | 04.09.2019 | 50.0556 | 8.7788 | | x |
| S_147 | damaged | Heusenstamm | 04.09.2019 | 50.056 | 8.78 | dSouth | |
| S_148 | healthy | Heusenstamm | 04.09.2019 | 50.056 | 8.78 | hSouth | |
| S_149 | damaged | Heusenstamm | 04.09.2019 | 50.0557 | 8.779 | dSouth | |
| S_150 | healthy | Heusenstamm | 04.09.2019 | 50.0557 | 8.779 | hSouth | |
| S_151 | damaged | Heusenstamm | 04.09.2019 | 50.0557 | 8.779 | dSouth | |
| S_152 | healthy | Heusenstamm | 04.09.2019 | 50.0557 | 8.779 | hSouth | |
| S_153 | damaged | Heusenstamm | 04.09.2019 | 50.0552 | 8.7797 | dSouth | |
| S_154 | healthy | Heusenstamm | 04.09.2019 | 50.0552 | 8.7797 | hSouth | |
| S_155 | damaged | Eschollbrücken | 06.09.2019 | 49.844862 | 8.6033987 | dSouth | |

| | | | | | | | |
|---|---|---|---|---|---|---|---|
| S_156 | healthy | Eschollbrücken | 06.09.2019 | 49.844862 | 8.6033987 | hSouth | |
| S_157 | damaged | Eschollbrücken | 06.09.2019 | 49.8451033 | 8.6033906 | dSouth | |
| S_158 | healthy | Eschollbrücken | 06.09.2019 | 49.8451033 | 8.6033906 | hSouth | |
| S_159 | damaged | Eschollbrücken | 06.09.2019 | 49.8452741 | 8.6933477 | dSouth | |
| S_160 | healthy | Eschollbrücken | 06.09.2019 | 49.8452741 | 8.6933477 | hSouth | |
| S_161 | damaged | Eschollbrücken | 06.09.2019 | 49.8464188 | 8.6058807 | dSouth | |
| S_162 | healthy | Eschollbrücken | 06.09.2019 | 49.8464188 | 8.6058807 | hSouth | |
| S_163 | damaged | Eschollbrücken | 06.09.2019 | 49.8461255 | 8.6064631 | | x |
| S_164 | healthy | Eschollbrücken | 06.09.2019 | 49.8461255 | 8.6064631 | | x |
| S_165 | damaged | Eschollbrücken | 06.09.2019 | 49.8381468 | 8.613406 | dSouth | |
| S_166 | healthy | Eschollbrücken | 06.09.2019 | 49.8381468 | 8.613406 | hSouth | |
| S_167 | damaged | Eschollbrücken | 06.09.2019 | 49.8371837 | 8.6154874 | dSouth | |
| S_168 | healthy | Eschollbrücken | 06.09.2019 | 49.8371837 | 8.6154874 | hSouth | |
| S_169 | damaged | Eschollbrücken | 06.09.2019 | 49.8368364 | 8.6172492 | | x |
| S_170 | healthy | Eschollbrücken | 06.09.2019 | 49.8368364 | 8.6172492 | | x |
| S_171 | damaged | Eschollbrücken | 06.09.2019 | 49.8364971 | 8.6193427 | dSouth | |
| S_172 | healthy | Eschollbrücken | 06.09.2019 | 49.8364971 | 8.6193427 | hSouth | |
| S_173 | damaged | Eschollbrücken | 06.09.2019 | 49.8366439 | 8.617927 | dSouth | |
| S_174 | healthy | Eschollbrücken | 06.09.2019 | 49.8366439 | 8.617927 | hSouth | |
| S_175 | damaged | Eschollbrücken | 06.09.2019 | 49.83549 | 8.6063276 | | x |
| S_176 | healthy | Eschollbrücken | 06.09.2019 | 49.83549 | 8.6063276 | | x |
| S_177 | damaged | Eschollbrücken | 06.09.2019 | 49.836002 | 8.6057516 | dSouth | |
| S_178 | healthy | Eschollbrücken | 06.09.2019 | 49.836002 | 8.6057516 | hSouth | |
| S_179 | damaged | Eschollbrücken | 06.09.2019 | 49.8364465 | 8.6051897 | | x |
| S_180 | healthy | Eschollbrücken | 06.09.2019 | 49.8364465 | 8.6051897 | | x |
| S_181 | damaged | Neu Isenburg | 09.09.2019 | 50.0643187 | 8.7090884 | dSouth | |
| S_182 | healthy | Neu Isenburg | 09.09.2019 | 50.0643187 | 8.7090884 | hSouth | |
| S_183 | damaged | Neu Isenburg | 09.09.2019 | 50.0650797 | 8.7098096 | dSouth | |
| S_184 | healthy | Neu Isenburg | 09.09.2019 | 50.0650797 | 8.7098096 | hSouth | |
| S_185 | damaged | Neu Isenburg | 09.09.2019 | 50.0659154 | 8.7102773 | dSouth | |
| S_186 | healthy | Neu Isenburg | 09.09.2019 | 50.0659154 | 8.7102773 | hSouth | |
| S_187 | damaged | Neu Isenburg | 09.09.2019 | 50.0659301 | 8.7100359 | dSouth | |
| S_188 | healthy | Neu Isenburg | 09.09.2019 | 50.0659301 | 8.7100359 | hSouth | |
| S_189 | damaged | Neu Isenburg | 09.09.2019 | 50.0686111 | 8.7121451 | dSouth | |
| S_190 | healthy | Neu Isenburg | 09.09.2019 | 50.0686111 | 8.7121451 | hSouth | |
| S_191 | damaged | Neu Isenburg | 09.09.2019 | 50.0691756 | 8.7123064 | dSouth | |
| S_192 | healthy | Neu Isenburg | 09.09.2019 | 50.0691756 | 8.7123064 | hSouth | |
| S_193 | damaged | Neu Isenburg | 09.09.2019 | 50.0710306 | 8.7127781 | dSouth | |

| | | | | | | | |
|---|---|---|---|---|---|---|---|
| S_194 | healthy | Neu Isenburg | 09.09.2019 | 50.0710306 | 8.7127781 | hSouth | |
| S_195 | damaged | Neu Isenburg | 09.09.2019 | 50.0716691 | 8.7111252 | dSouth | |
| S_196 | healthy | Neu Isenburg | 09.09.2019 | 50.0716691 | 8.7111252 | hSouth | |
| S_197 | damaged | Neu Isenburg | 09.09.2019 | 50.0715402 | 8.7110297 | dSouth | |
| S_198 | healthy | Neu Isenburg | 09.09.2019 | 50.0715402 | 8.7110297 | hSouth | |
| S_199 | damaged | Neu Isenburg | 09.09.2019 | 50.0710603 | 8.7121176 | dSouth | |
| S_200 | healthy | Neu Isenburg | 09.09.2019 | 50.0710603 | 8.7121176 | hSouth | |
| S_201 | damaged | Neu Isenburg | 09.09.2019 | 50.0681943 | 8.7052592 | dSouth | |
| S_202 | healthy | Neu Isenburg | 09.09.2019 | 50.0681943 | 8.7052592 | hSouth | |
| S_203 | damaged | Neu Isenburg | 09.09.2019 | 50.0671452 | 8.7042427 | dSouth | |
| S_204 | healthy | Neu Isenburg | 09.09.2019 | 50.0671452 | 8.7042427 | hSouth | |
| S_205 | damaged | Wegscheide | 10.09.2019 | 50.2076 | 9.4126 | | x |
| S_206 | healthy | Wegscheide | 10.09.2019 | 50.2076 | 9.4126 | | x |
| S_207 | damaged | Wegscheide | 10.09.2019 | 50.2076 | 9.4126 | dSouth | |
| S_208 | healthy | Wegscheide | 10.09.2019 | 50.2076 | 9.4126 | hSouth | |
| S_209 | damaged | Wegscheide | 10.09.2019 | 50.2149354 | 9.419513 | dSouth | |
| S_210 | healthy | Wegscheide | 10.09.2019 | 50.2149354 | 9.419513 | hSouth | |
| S_211 | damaged | Wegscheide | 10.09.2019 | 50.2114485 | 9.409662 | dSouth | |
| S_212 | healthy | Wegscheide | 10.09.2019 | 50.2114485 | 9.409662 | hSouth | |
| S_213 | damaged | Wegscheide | 10.09.2019 | 50.2114485 | 9.409662 | dSouth | |
| S_214 | healthy | Wegscheide | 10.09.2019 | 50.2114485 | 9.409662 | hSouth | |
| S_215 | damaged | Wegscheide | 10.09.2019 | 50.2109 | 9.4086 | dSouth | |
| S_216 | healthy | Wegscheide | 10.09.2019 | 50.2109 | 9.4086 | hSouth | |
| S_217 | damaged | Waldstadion | 10.09.2019 | 50.0546504 | 8.6626215 | dSouth | |
| S_218 | healthy | Waldstadion | 10.09.2019 | 50.0546504 | 8.6626215 | hSouth | |
| S_219 | damaged | Waldstadion | 10.09.2019 | 50.0555269 | 8.6618698 | dSouth | |
| S_220 | healthy | Waldstadion | 10.09.2019 | 50.0555269 | 8.6618698 | hSouth | |
| S_221 | damaged | Waldstadion | 10.09.2019 | 50.0568445 | 8.6610993 | dSouth | |
| S_222 | healthy | Waldstadion | 10.09.2019 | 50.0568445 | 8.6610993 | hSouth | |
| S_223 | damaged | Waldstadion | 10.09.2019 | 50.0572346 | 8.6619006 | dSouth | |
| S_224 | healthy | Waldstadion | 10.09.2019 | 50.0572346 | 8.6619006 | hSouth | |
| S_225 | damaged | Waldstadion | 10.09.2019 | 50.0573499 | 8.661843 | dSouth | |
| S_226 | healthy | Waldstadion | 10.09.2019 | 50.0573499 | 8.661843 | hSouth | |
| S_227 | damaged | Waldstadion | 10.09.2019 | 50.0589663 | 8.6609092 | dSouth | |
| S_228 | healthy | Waldstadion | 10.09.2019 | 50.0589663 | 8.6609092 | hSouth | |
| S_229 | damaged | Waldstadion | 10.09.2019 | 50.0587827 | 8.6609424 | dSouth | |
| S_230 | healthy | Waldstadion | 10.09.2019 | 50.0587827 | 8.6609424 | hSouth | |
| S_231 | damaged | Messel | 11.09.2019 | 49.9648055 | 8.7305273 | dSouth | |

| | | | | | | | |
|---|---|---|---|---|---|---|---|
| S_232 | healthy | Messel | 11.09.2019 | 49.9648055 | 8.7305273 | hSouth | |
| S_233 | damaged | Messel | 11.09.2019 | 49.9653231 | 8.7286833 | dSouth | |
| S_234 | healthy | Messel | 11.09.2019 | 49.9653231 | 8.7286833 | hSouth | |
| S_235 | damaged | Messel | 11.09.2019 | 49.9672891 | 8.7267089 | dSouth | |
| S_236 | healthy | Messel | 11.09.2019 | 49.9672891 | 8.7267089 | hSouth | |
| S_237 | damaged | Messel | 11.09.2019 | 49.9673648 | 8.726613 | dSouth | |
| S_238 | healthy | Messel | 11.09.2019 | 49.9673648 | 8.726613 | hSouth | |
| S_239 | damaged | Messel | 11.09.2019 | 49.9685641 | 8.7259826 | dSouth | |
| S_240 | healthy | Messel | 11.09.2019 | 49.9685641 | 8.7259826 | hSouth | |
| S_241 | damaged | Messel | 11.09.2019 | 49.9702459 | 8.7244602 | dSouth | |
| S_242 | healthy | Messel | 11.09.2019 | 49.9702459 | 8.7244602 | hSouth | |
| S_243 | damaged | Messel | 11.09.2019 | 49.9722449 | 8.7156846 | dSouth | |
| S_244 | healthy | Messel | 11.09.2019 | 49.9722449 | 8.7156846 | hSouth | |
| S_245 | damaged | Messel | 11.09.2019 | 49.9722449 | 8.7156806 | | x |
| S_246 | healthy | Messel | 11.09.2019 | 49.9722449 | 8.7156806 | | x |
| S_247 | damaged | Messel | 11.09.2019 | 49.9724476 | 8.7142302 | dSouth | |
| S_248 | healthy | Messel | 11.09.2019 | 49.9724476 | 8.7142302 | hSouth | |
| S_249 | damaged | Messel | 11.09.2019 | 49.9718971 | 8.7119104 | dSouth | |
| S_250 | healthy | Messel | 11.09.2019 | 49.9718971 | 8.7119104 | hSouth | |
| S_251 | damaged | Messel | 11.09.2019 | 49.9719081 | 8.7114799 | dSouth | |
| S_252 | healthy | Messel | 11.09.2019 | 49.9719081 | 8.7114799 | hSouth | |
| S_253 | damaged | Lorsch | 12.09.2019 | 49.6491886 | 8.5376088 | dSouth | |
| S_254 | healthy | Lorsch | 12.09.2019 | 49.6491886 | 8.5376088 | hSouth | |
| S_255 | damaged | Lorsch | 12.09.2019 | 49.6493532 | 8.5369721 | dSouth | |
| S_256 | healthy | Lorsch | 12.09.2019 | 49.6493532 | 8.5369721 | hSouth | |
| S_257 | damaged | Lorsch | 12.09.2019 | 49.6488428 | 8.5381241 | dSouth | |
| S_258 | healthy | Lorsch | 12.09.2019 | 49.6488428 | 8.5381241 | hSouth | |
| S_259 | damaged | Lorsch | 12.09.2019 | 49.6485083 | 8.5392349 | dSouth | |
| S_260 | healthy | Lorsch | 12.09.2019 | 49.6485083 | 8.5392349 | hSouth | |
| S_261 | damaged | Lorsch | 12.09.2019 | 49.6481419 | 8.5395785 | dSouth | |
| S_262 | healthy | Lorsch | 12.09.2019 | 49.6481419 | 8.5395785 | hSouth | |
| S_263 | damaged | Lorsch | 12.09.2019 | 49.6462656 | 8.5421035 | dSouth | |
| S_264 | healthy | Lorsch | 12.09.2019 | 49.6462656 | 8.5421035 | hSouth | |
| S_265 | damaged | Lorsch | 12.09.2019 | 49.6464364 | 8.5415305 | | x |
| S_266 | healthy | Lorsch | 12.09.2019 | 49.6464364 | 8.5415305 | | x |
| S_267 | damaged | Lorsch | 12.09.2019 | 49.6463205 | 8.5410504 | dSouth | |
| S_268 | healthy | Lorsch | 12.09.2019 | 49.6463205 | 8.5410504 | hSouth | |
| S_269 | damaged | Lorsch | 12.09.2019 | 49.6452786 | 8.5358174 | dSouth | |

| | | | | | | | |
|---|---|---|---|---|---|---|---|
| S_270 | healthy | Lorsch | 12.09.2019 | 49.6452786 | 8.5358174 | hSouth | |
| S_271 | damaged | Lorsch | 12.09.2019 | 49.6451675 | 8.5351421 | dSouth | |
| S_272 | healthy | Lorsch | 12.09.2019 | 49.6451675 | 8.5351421 | hSouth | |
| S_273 | damaged | Lorsch | 12.09.2019 | 49.649297 | 8.5362281 | | x |
| S_274 | healthy | Lorsch | 12.09.2019 | 49.649297 | 8.5362281 | | x |
| S_275 | damaged | Lorsch | 12.09.2019 | 49.6494678 | 8.5366794 | dSouth | |
| S_276 | healthy | Lorsch | 12.09.2019 | 49.6494678 | 8.5366794 | hSouth | |
| S_277 | damaged | Heusenstamm | 13.09.2019 | 50.0580508 | 8.7747529 | dSouth | |
| S_278 | healthy | Heusenstamm | 13.09.2019 | 50.0580508 | 8.7747529 | hSouth | |
| S_279 | damaged | Heusenstamm | 13.09.2019 | 50.0580635 | 8.7746208 | | x |
| S_280 | healthy | Heusenstamm | 13.09.2019 | 50.0580635 | 8.7746208 | | x |
| S_281 | damaged | Heusenstamm | 13.09.2019 | 50.057182 | 8.773846 | dSouth | |
| S_282 | healthy | Heusenstamm | 13.09.2019 | 50.057182 | 8.773846 | hSouth | |
| S_283 | damaged | Heusenstamm | 13.09.2019 | 50.0571046 | 8.7740264 | dSouth | |
| S_284 | healthy | Heusenstamm | 13.09.2019 | 50.0571046 | 8.7740264 | hSouth | |
| S_285 | damaged | Heusenstamm | 13.09.2019 | 50.0566422 | 8.7736616 | dSouth | |
| S_286 | healthy | Heusenstamm | 13.09.2019 | 50.0566422 | 8.7736616 | hSouth | |
| S_287 | damaged | Heusenstamm | 13.09.2019 | 50.0561296 | 8.7732438 | dSouth | |
| S_288 | healthy | Heusenstamm | 13.09.2019 | 50.0561296 | 8.7732438 | hSouth | |
| S_289 | damaged | Heusenstamm | 13.09.2019 | 50.0559955 | 8.7728465 | dSouth | |
| S_290 | healthy | Heusenstamm | 13.09.2019 | 50.0559955 | 8.7728465 | hSouth | |
| S_291 | damaged | Heusenstamm | 13.09.2019 | 50.055472 | 8.7729314 | dSouth | |
| S_292 | healthy | Heusenstamm | 13.09.2019 | 50.055472 | 8.7729314 | hSouth | |
| S_293 | damaged | Heusenstamm | 13.09.2019 | 50.055181 | 8.772228 | dSouth | |
| S_294 | healthy | Heusenstamm | 13.09.2019 | 50.055181 | 8.772228 | hSouth | |
| S_295 | damaged | Heusenstamm | 13.09.2019 | 50.0551293 | 8.7723349 | dSouth | |
| S_296 | healthy | Heusenstamm | 13.09.2019 | 50.0551293 | 8.7723349 | hSouth | |
| S_297 | damaged | Heusenstamm | 13.09.2019 | 50.0549188 | 8.7721502 | dSouth | |
| S_298 | healthy | Heusenstamm | 13.09.2019 | 50.0549188 | 8.7721502 | hSouth | |
| S_299 | damaged | Heusenstamm | 13.09.2019 | 50.0548086 | 8.7726598 | | X |
| S_300 | healthy | Heusenstamm | 13.09.2019 | 50.0548086 | 8.7726598 | | X |
| conf_001 | damaged | Schloßborn | 03.08.2020 | 50.1904623 | 8.373935 | | x |
| conf_002 | healthy | Schloßborn | 03.08.2020 | 50.1902617 | 8.3738026 | | X |
| conf_003 | damaged | Schloßborn | 03.08.2020 | 50.1887401 | 8.372701 | | |
| conf_004 | healthy | Schloßborn | 03.08.2020 | 50.1879031 | 8.3716613 | | X |
| conf_005 | damaged | Schloßborn | 03.08.2020 | 50.1865722 | 8.3710684 | | X |
| conf_006 | healthy | Schloßborn | 03.08.2020 | 50.18656127 | 8.3711265 | | X |
| conf_007 | damaged | Schloßborn | 03.08.2020 | 50.1857023 | 8.3708097 | | X |

| | | | | | | |
|---|---|---|---|---|---|---|
| conf_008 | healthy | Schloßborn | 03.08.2020 | 50.1851555 | 8.3712359 | X |
| conf_009 | damaged | Schloßborn | 03.08.2020 | 50.1852938 | 8.371852 | X |
| conf_010 | healthy | Schloßborn | 03.08.2020 | 50.1851807 | 8.3722412 | X |
| conf_011 | healthy | Schloßborn | 03.08.2020 | 50.1864421 | 8.3767229 | X |
| conf_012 | damaged | Schloßborn | 03.08.2020 | 50.1864287 | 8.3764222 | X |
| conf_013 | damaged | Ruppertshain | 04.08.2020 | 50.181629 | 8.4018675 | X |
| conf_014 | healthy | Ruppertshain | 04.08.2020 | 50.1815555 | 8.4018799 | X |
| conf_016 | healthy | Ruppertshain | 04.08.2020 | 50.1818417 | 8.4023975 | X |
| conf_017 | damaged | Ruppertshain | 04.08.2020 | 50.1846569 | 8.4022888 | X |
| conf_018 | healthy | Ruppertshain | 04.08.2020 | 50.1847183 | 8.40225810 | X |
| conf_019 | damaged | Ruppertshain | 04.08.2020 | 50.1867665 | 8.4029614 | X |
| conf_020 | healthy | Ruppertshain | 04.08.2020 | 50.1868466 | 8.4029997 | X |
| conf_021 | damaged | Ruppertshain | 04.08.2020 | 50.1868411 | 8.4040772 | X |
| conf_022 | healthy | Ruppertshain | 04.08.2020 | 50.1877796 | 8.4046797 | X |
| conf_023 | damaged | Ruppertshain | 04.08.2020 | 50.1829212 | 8.3987642 | X |
| conf_024 | healthy | Ruppertshain | 04.08.2020 | 50.182956 | 8.3987399 | X |
| conf_025 | damaged | Ruppertshain | 04.08.2020 | 50.1808246 | 8.3990728 | X |
| conf_026 | healthy | Ruppertshain | 04.08.2020 | 50.1806971 | 8.39899759 | X |
| conf_027 | damaged | Ruppertshain | 04.08.2020 | 50.179967 | 8.3994577 | X |
| conf_028 | healthy | Ruppertshain | 04.08.2020 | 50.1798082 | 8.399525 | X |
| conf_029 | damaged | Ruppertshain | 04.08.2020 | 50.1782612 | 8.3999872 | X |
| conf_030 | healthy | Ruppertshain | 04.08.2020 | 50.1782916 | 8.4002846 | X |
| conf_032 | healthy | Falkenstein | 05.08.2020 | 50.201454 | 8.4872785 | X |
| conf_033 | damaged | Falkenstein | 05.08.2020 | 50.2015038 | 8.4874409 | X |
| conf_034 | damaged | Falkenstein | 05.08.2020 | 50.201115 | 8.4876516 | X |
| conf_036 | healthy | Falkenstein | 05.08.2020 | 50.2018627 | 8.4875841 | X |
| conf_037 | damaged | Falkenstein | 05.08.2020 | 50.2011419 | 8.4892266 | x |
| conf_038 | healthy | Falkenstein | 05.08.2020 | 50.2012289 | 8.489425 | x |
| conf_039 | healthy | Falkenstein | 05.08.2020 | 50.2013307 | 8.4905266 | |
| conf_040 | damaged | Falkenstein | 05.08.2020 | 50.2004517 | 8.4905881 | x |
| conf_041 | healthy | Falkenstein | 05.08.2020 | 50.2008376 | 8.4909408 | x |
| conf_042 | damaged | Falkenstein | 05.08.2020 | 50.20009229 | 8.4909295 | x |
| conf_043 | damaged | Falkenstein | 05.08.2020 | 50.1997086 | 8.4828883 | x |
| conf_045 | healthy | Falkenstein | 05.08.2020 | 50.1992717 | 8.480502 | x |
| conf_046 | damaged | Falkenstein | 05.08.2020 | 50.1995271 | 8.4806157 | x |
| conf_047 | healthy | Falkenstein | 05.08.2020 | 50.1995032 | 8.4806215 | x |
| conf_050 | damaged | Rote Mühle | 07.08.2020 | 50.1612598 | 8.452304 | x |
| conf_051 | healthy | Rote Mühle | 07.08.2020 | 50.1610649 | 8.4526065 | x |

| | | | | | | | |
|---|---|---|---|---|---|---|---|
| conf_052 | damaged | Rote Mühle | 07.08.2020 | 50.1611651 | 8.4520887 | | x |
| conf_053 | healthy | Rote Mühle | 07.08.2020 | 50.1611113 | 8.4520672 | | x |
| conf_054 | damaged | Rote Mühle | 07.08.2020 | 50.1611776 | 8.4511498 | | x |
| conf_055 | healthy | Rote Mühle | 07.08.2020 | 50.1611331 | 8.450964 | | x |
| conf_056 | damaged | Rote Mühle | 07.08.2020 | 50.1612598 | 8.4508452 | | x |
| conf_057 | healthy | Rote Mühle | 07.08.2020 | 50.1614085 | 8.4509926 | | x |
| conf_058 | damaged | Rote Mühle | 07.08.2020 | 50.1617899 | 8.4504762 | | x |
| conf_059 | healthy | Rote Mühle | 07.08.2020 | 50.1617076 | 8.4505298 | | x |

**Suppl. Table 2**. Results from Mann-Whitney U-test on difference between damaged and healthy trees for various parameters.

| Parameter | Mann-Whitney U | p |
|---|---|---|
| trunk circumference | 10718 | 0.48 |
| tree height | 9874 | 0.27 |
| canopy closure | 10820 | 0.56 |
| competition index | 10809 | 0.57 |
| dried leaves | 202 | **<0.001** |
| leaf loss | 352 | **<0.001** |

**Suppl. Fig.1**. Climate change dynamics. Mean change of sampling site values along PCA axis 1 (temperature, red) and PCA axis 2 (precipitation, blue) among decades.

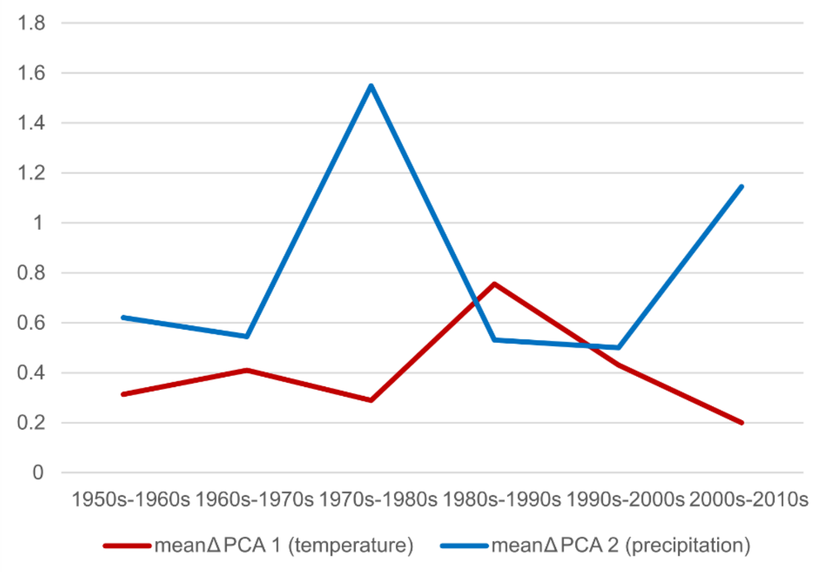


**Suppl. Fig. 2**. Plot of "relative Leaf Area Index" change relative to 2014 values against "cumulated evatransporation potential" change during the growth season relative to 2014 values for all 1 x 1 km plots encompassing the 27 sampling sites. The overall correlation is r = 0.695.

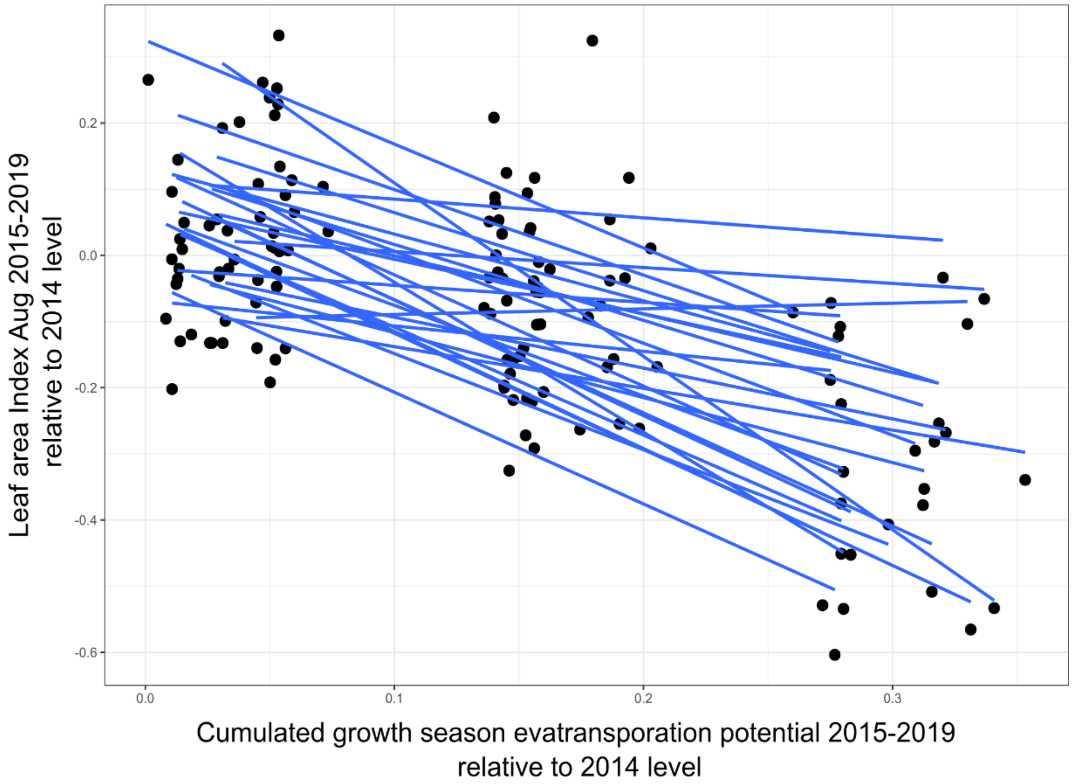

**Suppl. Fig. 3**. Distribution of pairwise distances between the paired trees.

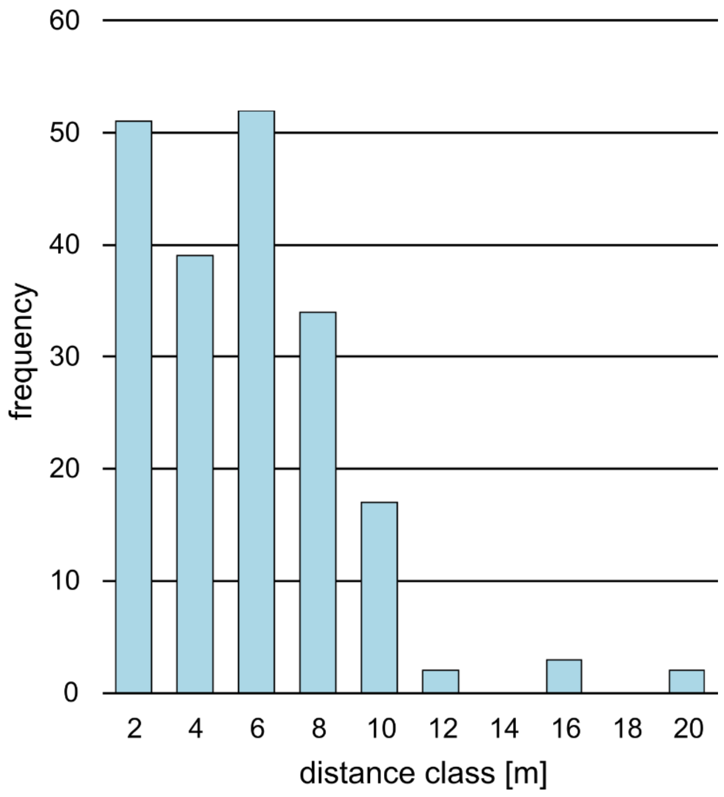

**Suppl. Fig. 4**. Exemplary pictures of damaged and healthy beech tree pairs from several sampling sites.

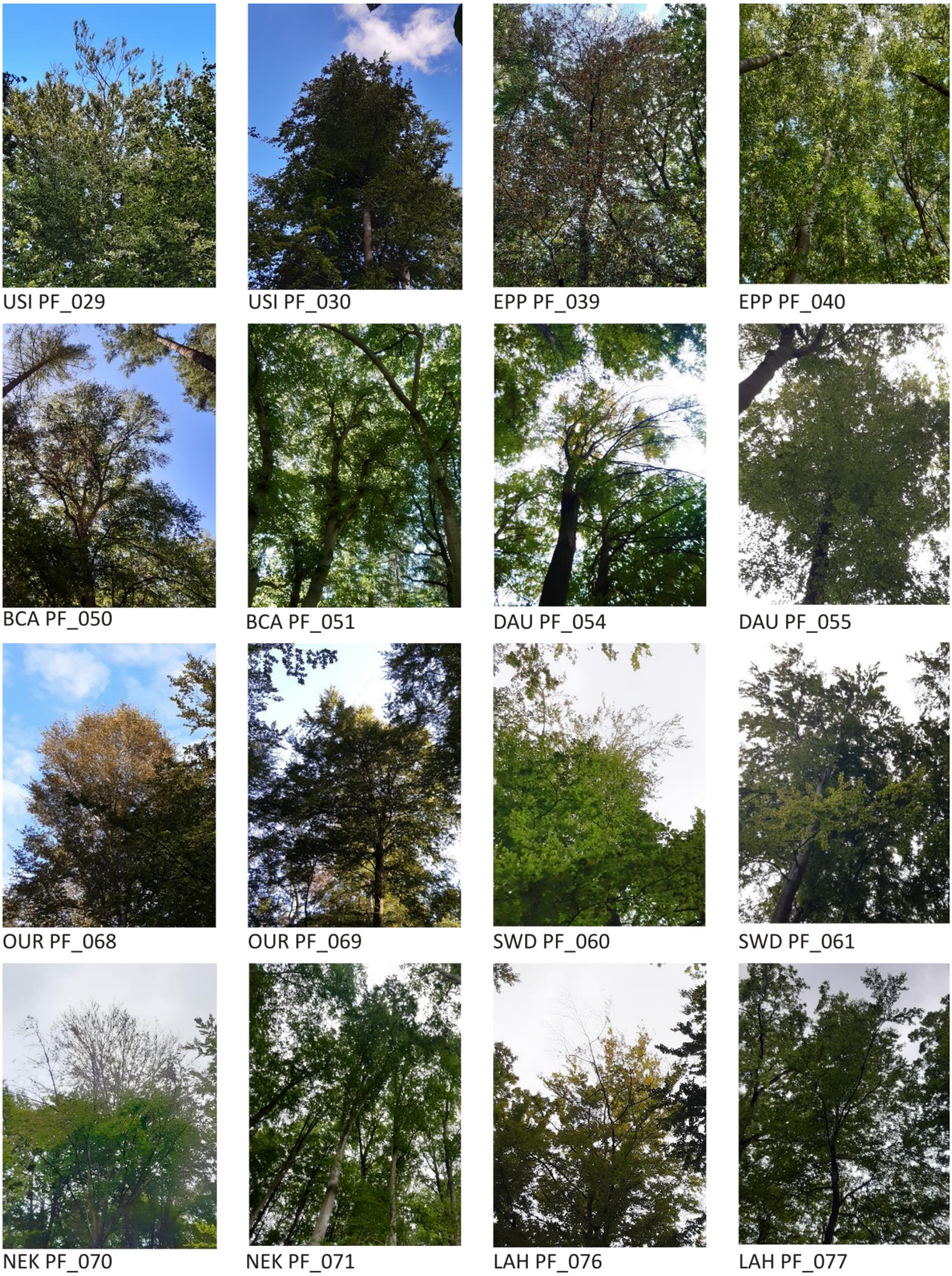

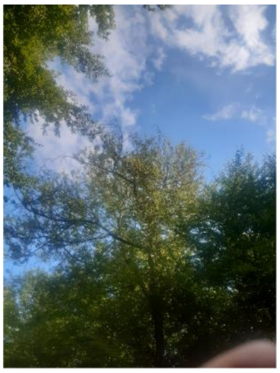
SLB PF_095

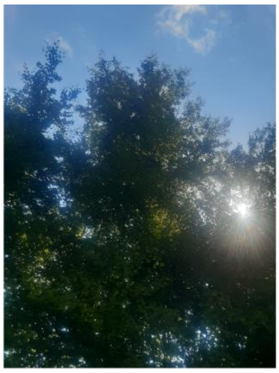
SLB PF_096

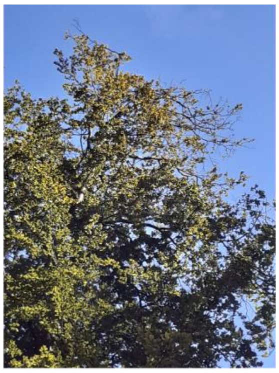
THT PF_089

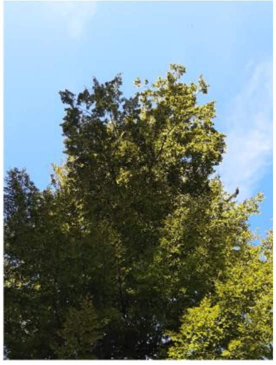
THT PF_090

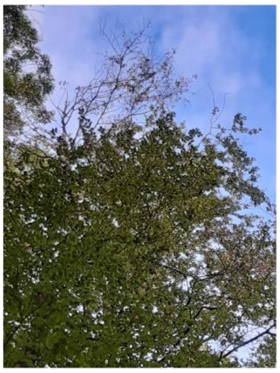
WEH PF_110

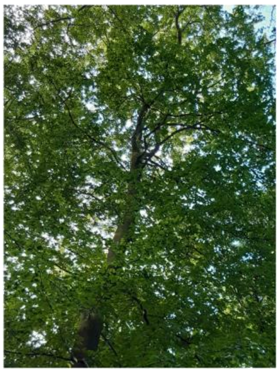
WEH PF_111

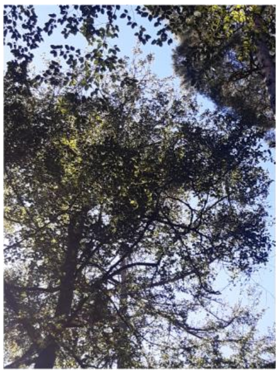
OBH PF_019

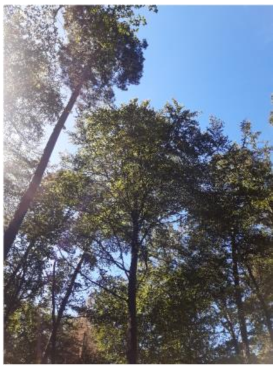
OBH PF_020

**Suppl. Fig. 5**. Genome-wide $F_{ST}$ distributions in 1 kb windows. Comparisons between all pools. Please note that the standard deviation included zero in all cases.

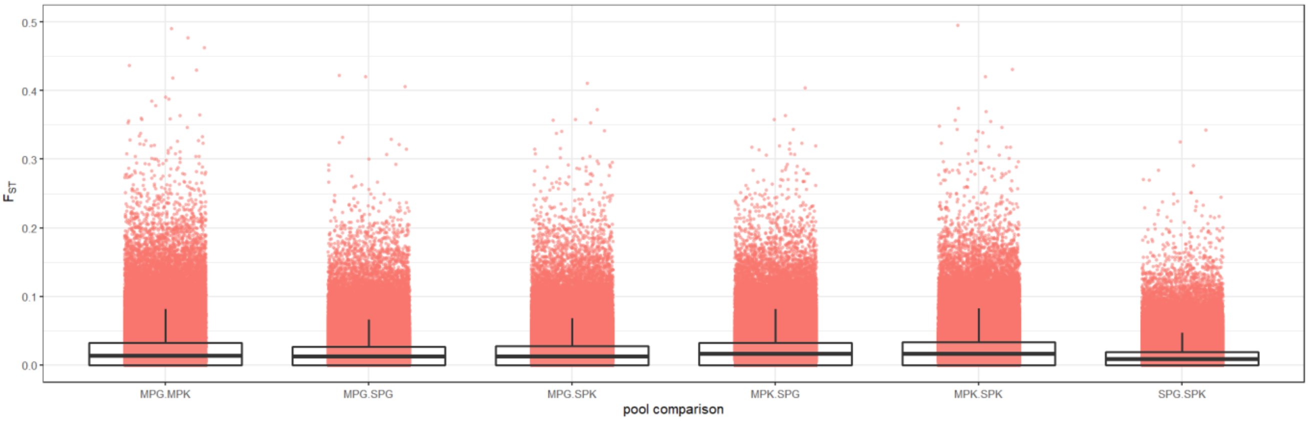

**Suppl. Fig. 6**. Genomic similarity among individuals within and among phenotypic classes. Permutation ANOSIM shows that the differences are not significant (p = 0.76).

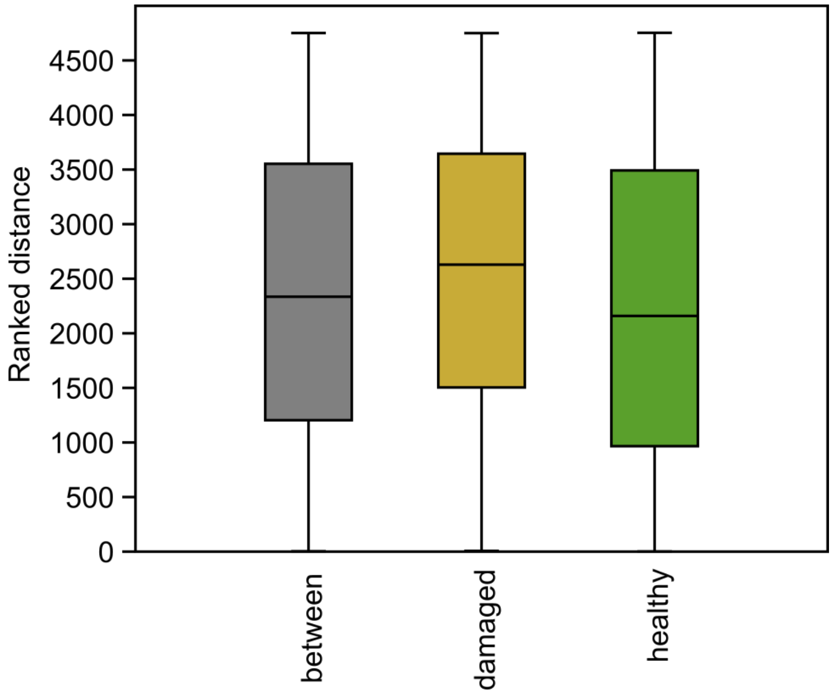

**Suppl. Fig. 7**. A) Manhattan plot of uncorrected p values from CMH test and B) corresponding QQ-plot. SNPs on different chromosomes are alternatingly coloured blue and black.

A)

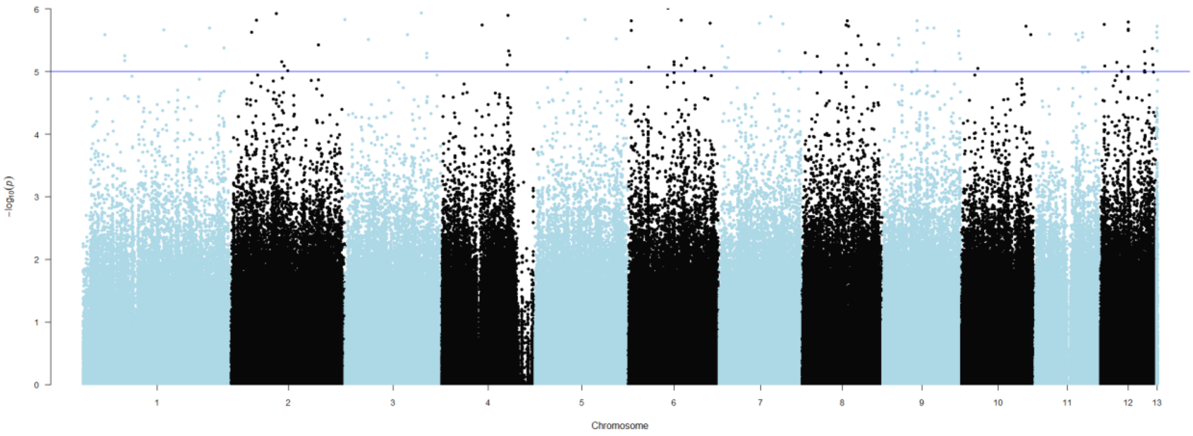


B)

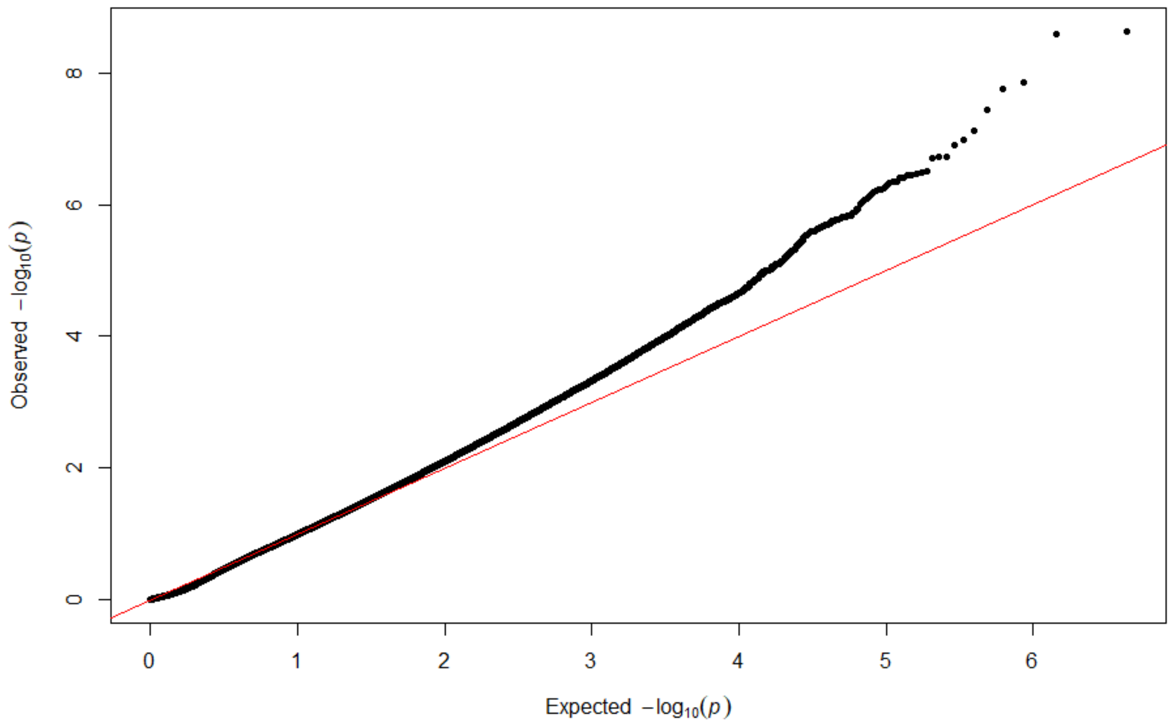

**Suppl. Fig. 8**. Increase of LDA prediction success with the number of loci involved. Loci were added according to their decreasing contribution in the final analysis.

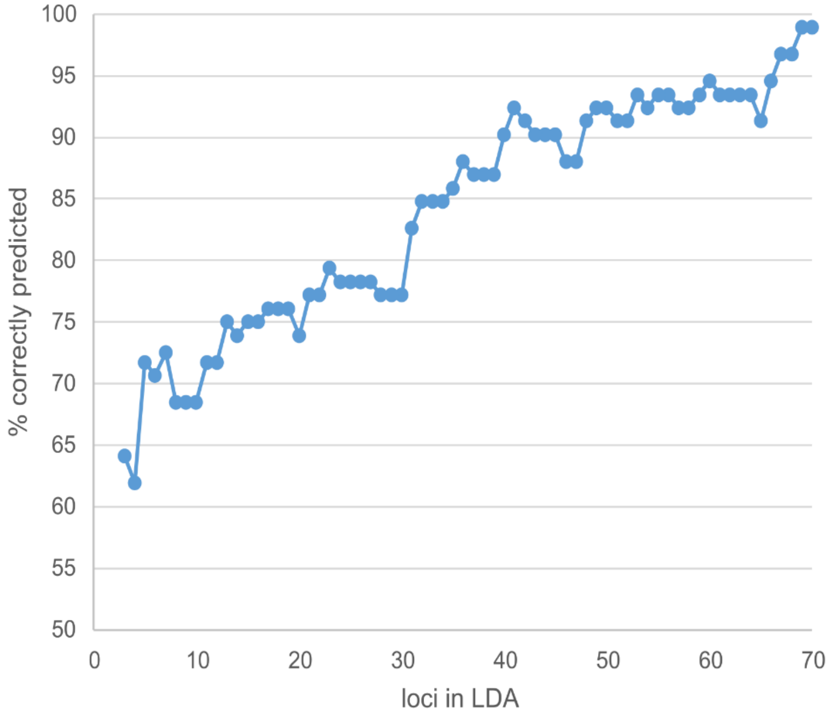

**Suppl. Fig. 9**. Comparison of observed heterozygosity between lower and upper half of predictive values “healthy” in DA. The medians are not significantly different (Mann-Whitney U = 293.5, p same median = 0.72).

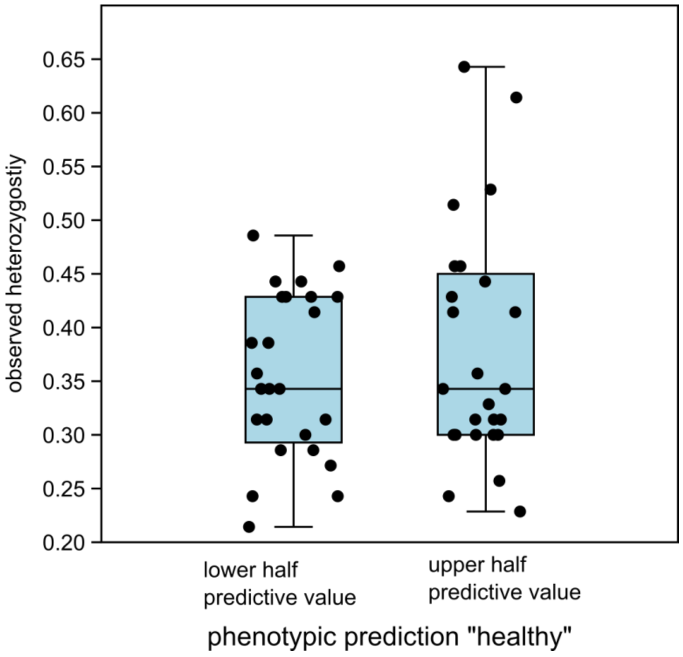


**Suppl. Fig.10**. Histogram of LDA results. Individuals, where the predicted phenotype did not match the observed phenotype are shown in grey, individuals with matching observed/predicted phenotype in green (healthy) or in ochre (damaged).

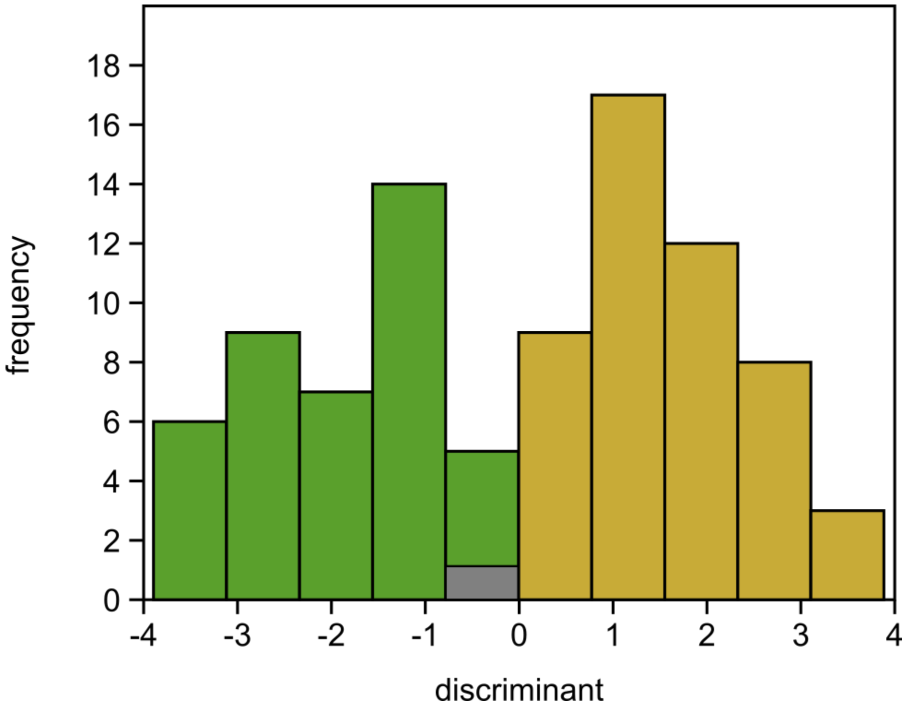

**Suppl. Table 3**. List of genes with significant SNPs. Functional annotations and relation to drought phenotype are given whenever available.

| ***F. sylvatica* gene ID** | **Best BLAST hit ID** | **BLAST hit protein name** | **taxon** | **UniProt ID** | **UniProt protein name** | **function (UniProt)** | **citation function** | **relation to drought phenotype** | **citation** |
|---|---|---|---|---|---|---|---|---|---|
| 1.g3851.t1 | KUM50718.1 | hypothetical protein ABT39_MTgene562 | *Picea glauca* | none | - | - | - | - | - |
| 10.g3914.t1 | XP_023883481.1 | uncharacterized protein LOC111995782 isoform X1 | *Quercus suber* | none | - | - | - | - | - |
| 11.g2467.t1 | XP_023923514.1 | exosome complex exonuclease RRP46 homolog isoform X1 | *Quercus suber* | EXOS5_ORYSJ | Exosome complex exonuclease RRP46 homolog | mRNA degradation | Xiang, D., Yang, H., Venglat, P., Cao, Y., Wen, R., Ren, M., ... & Weijers, D. (2011). POPCORN functions in the auxin pathway to regulate embryonic body plan and meristem organization in Arabidopsis. The Plant Cell, 23(12), 4348-4367. | - | - |
| 11.g2832.t1 | XP_030949821.1 | WD repeat-containing protein PCN-like | *Quercus lobata* | PCN_ARATH | WD repeat-containing protein PCN | Involved in auxin signalling pathway. Required for embryo development and meristem organization. Functions in the auxin pathway, integrating auxin signalling in the organization | - | drought stress response | Park, S. R., Hwang, J., & Kim, M. (2020). The Arabidopsis WDR55 is positively involved in ABA-mediated |

| | | | | | | | | | |
|---|---|---|---|---|---|---|---|---|---|
| | | | | | | and maintenance of the shoot apical meristem (SAM) and root apical meristem (RAM). | | | drought tolerance response. Plant Biotechnology Reports, 1-12. |
| 12.g1695.t1 | XP_030973623.1 | uncharacterized protein LOC115993791 | *Quercus lobata* | F4I5S1_ARATH | PB1 domain-containing protein tyrosine kinase | not well characterised | - | - | - |
| 2.g4736.t1 | KAF3975221.1 | hypothetical protein CMV_001513 | *Castanea mollissima* | none | - | - | - | - | - |
| 3.g3590.t1 | XP_030942104.1 | uncharacterized protein LOC115967180 | *Quercus lobata* | none | - | - | - | - | - |
| 4.g3980.t1 | XP_030950630.1 | cytokinin dehydrogenase 5 | *Quercus lobata* | CKX1_ARATH | Cytokinin dehydrogenase 1 | Catalyzes the oxidation of cytokinins, a family of N6-substituted adenine derivatives that are plant hormones, where the substituent is an isopentenyl group. | Werner, T., Motyka, V., Laucou, V., Smets, R., Van Onckelen, H., & Schmülling, T. (2003). Cytokinin-deficient transgenic Arabidopsis plants show multiple developmental alterations indicating opposite functions of cytokinins in the regulation of shoot and root meristem activity. The Plant Cell, 15(11), 2532- | drought stress response | Emery, R. J., & Kisiala, A. (2020). The Roles of Cytokinins in Plants and Their Response to Environmental Stimuli. Li, W., Herrera-Estrella, L., & Tran, L. S. P. (2016). The Yin–Yang of cytokinin homeostasis and drought acclimation/adaptation. Trends in Plant Science, |

| | | | | | | | | | |
|---|---|---|---|---|---|---|---|---|---|
| | | | | | | | 2550. | | 21(7), 548-550. |
| 5.g1807.t1 | XP_030939294.1 | guanosine nucleotide diphosphate dissociation inhibitor 2 | *Quercus lobata* | GDI2_ARATH | Guanosine nucleotide diphosphate dissociation inhibitor 2 | Regulates the GDP/GTP exchange reaction of most RAB proteins by inhibiting the dissociation of GDP from them, and the subsequent binding of GTP. | Ueda, T., Yoshizumi, T., Anai, T., Matsui, M., Uchimiya, H., & Nakano, A. (1998). AtGDI2, a novel Arabidopsis gene encoding a Rab GDP dissociation inhibitor. Gene, 206(1), 137-143. | environmental stress response | Carvalho, B. M., Viana, A. P., dos Santos, P. H. D., Generoso, A. L., Corrêa, C. C. G., Silveira, V., ... & Santos, E. A. (2019). Proteome of resistant and susceptible Passiflora species in the interaction with cowpea aphid-borne mosaic virus reveals distinct responses to pathogenesis. Euphytica, 215(10), 167. |
| 6.g2227.t1 | THG21676.1 | hypothetical protein TEA_000305 | *Camellia sinensis* var. *sinensis* | NDUS7_ARATH | NADH dehydrogenase [ubiquinone] iron-sulfur protein 7, mitochondrial | Core subunit of the mitochondrial membrane respiratory chain NADH dehydrogenase (Complex I) that is believed to belong to the minimal assembly required for catalysis. Complex I functions in the transfer of electrons from NADH to the respiratory chain. The immediate electron acceptor for the enzyme is believed to be ubiquinone | | drought stress response | Zhang, S., Zhang, L., Zhou, K., Li, Y., & Zhao, Z. (2017). Changes in protein profile of Platycladus orientalis (L.) roots and leaves in response to drought stress. Tree Genetics & Genomes, 13(4), 76. |

| | | | | | | | | | |
|---|---|---|---|---|---|---|---|---|---|
| 6.g2921.t1 | KAB1222126.1 | Ribonuclease H2 subunit C | *Morella rubra* | RNH2A_ARATH | Ribonuclease H2 subunit A | Catalytic subunit of RNase HII, an endonuclease that specifically degrades the RNA of RNA:DNA hybrids. Participates in DNA replication, possibly by mediating the removal of lagging-strand Okazaki fragment RNA primers during DNA replication. Mediates the excision of single ribonucleotides from DNA:RNA duplexes | - | - | - |
| 7.g177.t1 | XP_018842121.1 | PREDICTED: tubby-like F-box protein 8 isoform X1 | *Juglans regia* | TLP10_ARATH | Tubby-like F-box protein 10 | Component of SCF(ASK-cullin-F-box) E3 ubiquitin ligase complexes, which may mediate the ubiquitination and subsequent proteasomal degradation of target proteins | - | drought stress response | Xu, J., Xing, S., Sun, Q., Zhan, C., Liu, X., Zhang, S., & Wang, X. (2019). The expression of a tubby-like protein from Malus domestica (Md TLP7) enhances abiotic stress tolerance in Arabidopsis. BMC plant biology, 19(1), 1-8. |
| 7.g1655.t1 | XP_023909357.1 | histone deacetylase 6 | *Quercus suber* | HDA6 | Histone deacetylase 6 | Responsible for the deacetylation of lysine residues on the N-terminal part of the core histones (H2A, H2B, H3 and H4). Histone deacetylation gives a tag for epigenetic repression and plays an important role in transcriptional regulation, cell | "Identification of Arabidopsis histone deacetylase HDA6 mutants that affect transgene expression."<br><br>Murfett J., Wang X.-J., Hagen G., | drought stress | Zheng, Y., Ding, Y., Sun, X., Xie, S., Wang, D., Liu, X., ... & Zhou, D. X. (2016). Histone deacetylase HDA9 negatively regulates salt and drought stress |

| | | | | | | | | | |
|---|---|---|---|---|---|---|---|---|---|
| | | | | | | cycle progression and developmental events. | Guilfoyle T.J.<br>Plant Cell 13:1047-1061(2001) | | responsiveness in Arabidopsis. Journal of experimental botany, 67(6), 1703-1713. |
| 7.g2350.t1 | KAF3966828.1 | hypothetical protein CMV_009102 | *Castanea mollissima* | PRK4_ARATH | Pollen receptor-like kinase 4 | Receptor-like kinase involved in the control of pollen germination and pollen tube polar growth. Can phosphorylate ROPGEF1 in vitro | Chang, F., Gu, Y., Ma, H., & Yang, Z. (2013). AtPRK2 promotes ROP1 activation via RopGEFs in the control of polarized pollen tube growth. Molecular plant, 6(4), 1187-1201. | environmental stress response | Guo, J., Dong, X., Li, Y., & Wang, B. (2020). NaCl treatment markedly enhanced pollen viability and pollen preservation time of euhalophyte Suaeda salsa via up regulation of pollen development-related genes. Journal of plant research, 133(1), 57-71. |
| 7.g3617.t1 | XP_023924613.1 | protein LIGHT-DEPENDENT SHORT HYPOCOTYLS 10-like isoform X1 | *Quercus suber* | LSH4_ARATH | Protein LIGHT-DEPENDENT SHORT HYPOCOTYLS 4 | Probable transcription regulator that acts as a developmental regulator by promoting cell growth in response to light. May suppress organ differentiation in the boundary region | Takeda, S., Hanano, K., Kariya, A., Shimizu, S., Zhao, L., Matsui, M., ... & Aida, M. (2011). CUP-SHAPED COTYLEDON1 transcription factor activates the expression of LSH4 and LSH3, two members of the ALOG gene family, in | - | |

| | | | | | | | | | |
|---|---|---|---|---|---|---|---|---|---|
| | | | | | | | shoot organ boundary cells. The Plant Journal, 66(6), 1066-1077. | | |
| 7.g3816.t1 | XP_018811587.1 | ethylene-responsive transcription factor 12-like | *Juglans regia* | TAFCL_ARATH | Transcription initiation factor TFIID subunit 12b | TAFs are components of the transcription factor IID (TFIID) complex that is essential for mediating regulation of RNA polymerase transcription. Required for the expression of a subset of ethylene-responsive genes (By similarity). Involved in the negative regulation of cytokinin sensitivity | - | - | - |
| 7.g552.t1 | No hits found | | - | - | - | - | - | - | - |
| 8.g3494.t1 | XP_022750437.1 | V-type proton ATPase subunit C | *Durio zibethinus* | VATC_ARATH | V-type proton ATPase subunit C | Subunit of the peripheral V1 complex of vacuolar ATPase. Subunit C is necessary for the assembly of the catalytic sector of the enzyme and is likely to have a specific function in its catalytic activity. V-ATPase is responsible for acidifying a variety of intracellular compartments in eukaryotic cells | - | drought stress response | Kausar, R., Arshad, M., Shahzad, A., & Komatsu, S. (2013). Proteomics analysis of sensitive and tolerant barley genotypes under drought stress. Amino Acids, 44(2), 345-359. Li, J., Jia, H., Han, X., Zhang, J., Sun, P., Lu, M., & Hu, J. (2016). Selection of reliable reference genes for gene expression analysis under abiotic |

| | | | | | | | | | |
|---|---|---|---|---|---|---|---|---|---|
| | | | | | | | | | stresses in the desert biomass willow, *Salix psammophila*. Frontiers in plant science, 7, 1505. |
| 9.g3080.t1 | XP_030929046.1 | probable inactive purple acid phosphatase 29 | *Quercus lobata* | PPA14_ARATH | Probable inactive purple acid phosphatase 14 | - | - | drought stress response | Street, N. R., Skogström, O., Sjödin, A., Tucker, J., Rodríguez-Acosta, M., Nilsson, P., ... & Taylor, G. (2006). The genetics and genomics of the drought response in Populus. The Plant Journal, 48(3), 321-341. Prinsi, B., Negri, A. S., Failla, O., Scienza, A., & Espen, L. (2018). Root proteomic and metabolic analyses reveal specific responses to drought stress in differently tolerant grapevine rootstocks. BMC plant biology, 18(1), 126. |
| 9.g4504.t1 | KAF3976808.1 | hypothetical protein CMV_000062 | *Castanea mollissi* | TBL33_ARATH | Protein trichome birefring | Probable xylan acetyltransferase that plays a role in xylan acetylation and | Yuan, Y., Teng, Q., Zhong, R., Haghighat, M., | drought stress response | Shuai, P., Liang, D., Zhang, Z., Yin, W., & Xia, X. (2013). |

| | | | | | | | | | |
|---|---|---|---|---|---|---|---|---|---|
| | | | *ma* | | ence-like 33 | normal deposition of secondary cell walls | Richardson, E. A., & Ye, Z. H. (2016). Mutations of Arabidopsis TBL32 and TBL33 affect xylan acetylation and secondary wall deposition. PLoS One, 11(1), e0146460. | | Identification of drought-responsive and novel Populus trichocarpamicroRNAs by high-throughput sequencing and their targets using degradome analysis. Bmc Genomics, 14(1), 233. |

**Suppl. Table** 4. List of genes closest to significant SNPs. Functional annotations and relation to drought phenotype are given whenever available.

| ***F. sylvatica* gene ID** | **Best BLAST hit ID** | **BLAST hit protein name** | **taxon** | **UniProt ID** | **UniProt protein name** | **function (UniProt)** | **citation function** | **relation to drought phenotype** | **citation** |
|---|---|---|---|---|---|---|---|---|---|
| Backbone_621.g33 | KAF3957547 | hypothetical protein CMV_017449 | *Castanea mollissima* | | | | | | |
| Backbone_621.g49 | XP_030932040 | pathogen-related protein | *Quercus lobata* | | | | | | |
| 1.g1111 | KAF3976566 | hypothetical protein CMV_000247 | *Castanea mollissima* | ACBP4_ARATH | Acyl-CoA-binding domain-containing protein 4 | Binds medium- and long-chain acyl-CoA esters with very high affinity. Can interact in vitro with oleoyl-CoA, barely with palmitoyl-CoA, but not with arachidonyl-CoA. | Leung, K. C., Li, H. Y., Mishra, G., & Chye, M. L. (2005). ACBP4 and ACBP5, novel Arabidopsis acyl-CoA-binding proteins with kelch motifs that bind oleoyl-CoA. Plant molecular biology, 55(2), 297-309. | environmental stress response | Du, Z. Y., Arias, T., Meng, W., & Chye, M. L. (2016). Plant acyl-CoA-binding proteins: an emerging family involved in plant development and stress responses. Progress in Lipid Research, 63, 165-181. |
| 1.g2051 | XP_031256586 | aspartic proteinase nepenthesin-1-like | *Pistacia vera* | | | | | drought stress response | Prinsi, B., Negri, A. S., Failla, O., Scienza, A., & Espen, L. (2018). Root proteomic and metabolic analyses reveal specific responses to drought stress in differently |

| | | | | | | | | | |
|---|---|---|---|---|---|---|---|---|---|
| | | | | | | | | | tolerant grapevine rootstocks. BMC plant biology, 18(1), 126. |
| 1.g5025 | No hits found | | | | | | | | |
| 1.g6250 | XP_030925267 | BTB/POZ domain and ankyrin repeat-containing protein NOOT2 | *Quercus lobata* | NPR5_ARATH | Regulatory protein NPR5 | May act as a substrate-specific adapter of an E3 ubiquitin-protein ligase complex (CUL3-RBX1-BTB) which mediates the ubiquitination and subsequent proteasomal degradation of target proteins | Hepworth, S. R., Zhang, Y., McKim, S., Li, X., & Haughn, G. W. (2005). BLADE-ON-PETIOLE–dependent signaling controls leaf and floral patterning in Arabidopsis. The Plant Cell, 17(5), 1434-1448. | | |
| 1.g7050 | PQQ17864 | insulin-degrading enzyme-like 1 peroxisomal | *Prunus yedoensis* | IDE1_ARATH | insulin-degrading enzyme-like 1 peroxisomal | Peptidase that might be involved in pathogen or wound response. | Lingard, M. J., & Bartel, B. (2009). Arabidopsis LON2 is necessary for peroxisomal function and sustained matrix protein import. Plant physiology, 151(3), 1354-1365. | | |

| | | | | | | | | | |
|---|---|---|---|---|---|---|---|---|---|
| 2.g1138 | XP_018818502 | PREDICTED: uncharacterized protein LOC108989373 isoform X3 | *Juglans regia* | | | | | | |
| 2.g1372 | KAF3967401 | hypothetical protein CMV_008608 | *Castanea mollissima* | | | | | | |
| 2.g1982 | No hits found | | | | | | | | |
| 2.g4924 | TKY55186 | Spermine synthase | *Spatholobus suberectus* | SPSY_ARATH | Spermine synthase | | | drought stress response | Yamaguchi, K., Takahashi, Y., Berberich, T., Imai, A., Takahashi, T., Michael, A. J., & Kusano, T. (2007). A protective role for the polyamine spermine against drought stress in Arabidopsis. Biochemical and biophysical research communications, 352(2), 486-490. |
| 3.g1427 | XP_030940223 | uncharacterized protein LOC115965177 | *Quercus lobata* | | | | | | |
| 3.g4686 | KAF3964003 | hypothetical protein CMV_011671 | *Castanea mollissima* | | | | | | |
| 4.g1310 | KAF3954505 | hypothetical protein CMV_020159 | *Castanea mollissima* | Q9C6S8_ARATH | Pre-mRNA-splicing factor of RES complex protein | | | drought stress response | Qi, Y., Yao, X., Zhao, D., & Lu, L. (2018). Overexpression of SbSKIP, a pre-mRNA splicing factor from |

| | | | | | | | | | |
|---|---|---|---|---|---|---|---|---|---|
| | | | | | | | | | Sorghum bicolor, enhances root growth and drought tolerance in Petunia hybrida. Scientia Horticulturae, 240, 281-287. |
| 4.g3868 | KAF3971069 | hypothetical protein CMV_005309 | *Castanea mollissima* | | | | | | |
| 4.g3915 | KAF3976838 | hypothetical protein CMV_000012 | *Castanea mollissima* | | | | | | |
| 5.g2789 | XP_023898567 | uncharacterized protein LOC112010459 isoform X2 | *Quercus suber* | Q9M089_ARATH | Keratin-associated protein (DUF1218) | | | drought stress response | Xu, J., Yuan, Y., Xu, Y., Zhang, G., Guo, X., Wu, F., ... & Tang, Q. (2014). Identification of candidate genes for drought tolerance by whole-genome resequencing in maize. BMC Plant Biology, 14(1), 83. |
| 5.g3716 | XP_023870312 | dynamin-related protein 5A | *Quercus suber* | DRP5A_ARATH | Dynamin-related protein 5A | Probable microtubule-associated force-producing protein that is targeted to the forming cell plate during cytokinesis. May play a role in cell division | Miyagishima, S. Y., Kuwayama, H., Urushihara, H., & Nakanishi, H. (2008). Evolutionary linkage between eukaryotic cytokinesis and | drought stress response | Ren, Z., Zhang, D., Cao, L., Zhang, W., Zheng, H., Liu, Z., ... & Su, H. (2020). Functions and regulatory framework of ZmNST3 in maize under lodging and drought stress. Plant, Cell & Environment, 43(9), 2272-2286. |

| | | | | | | | | | |
|---|---|---|---|---|---|---|---|---|---|
| | | | | | | | chloroplast division by dynamin proteins. Proceedings of the National Academy of Sciences, 105(39), 15202-15207. | | |
| 5.g4697 | XP_023901205 | cytochrome b561 and DOMON domain-containing protein At4g12980-like | *Quercus suber* | | | | | | |
| 6.g180 | KNA23105 | hypothetical protein SOVF_026910 | *Spinacia oleracea* | | | | | | |
| 6.g3229 | XP_015867595 | transcription initiation factor TFIID subunit 11 | *Ziziphus jujuba* | TA14B_ARATH | Transcription initiation factor TFIID | Negative regulator of flowering controlling the H4K5 acetylation levels in the FLC and FT chromatin. Positively regulates FLC expression. | Zacharaki, V., Benhamed, M., Poulios, S., Latrasse, D., Papoutsoglou, P., Delarue, M., & Vlachonasios, K. E. (2012). The Arabidopsis ortholog of the YEATS domain containing protein YAF9a regulates flowering by | drought stress response | Parvathi, M. S., Nataraja, K. N., Reddy, Y. N., Naika, M. B., & Gowda, M. C. (2019). Transcriptome analysis of finger millet (Eleusine coracana (L.) Gaertn.) reveals unique drought responsive genes. Journal of genetics, 98(2), 46. |

| | | | | | | | | | |
|---|---|---|---|---|---|---|---|---|---|
| | | | | | | | controlling H4 acetylation levels at the FLC locus. Plant science, 196, 44-52. | | |
| 6.g823 | No hits found | | | | | | | | |
| 7.g1655 | XP_023909357 | histone deacetylase 6 | *Quercus suber* | HDA6_ARATH | Histone deacetylase 6 | Responsible for the deacetylation of lysine residues on the N-terminal part of the core histones (H2A, H2B, H3 and H4). | | drought stress response | Kim, J. M., Sasaki, T., Ueda, M., Sako, K., & Seki, M. (2015). Chromatin changes in response to drought, salinity, heat, and cold stresses in plants. Frontiers in plant science, 6, 114. |
| 7.g2958 | KAF3966011 | hypothetical protein CMV_009858 | *Castanea mollissima* | F4I6B2_ARATH | SIT4 phosphatase-associated family protein | | | | |
| 7.g3483 | XP_023874962 | uncharacterized protein LOC111987473 | *Quercus suber* | | | | | | |
| 7.g3669 | XP_023883887 | protein trichome birefringence-like 38 | *Quercus suber* | TBL38_ARATH | protein trichome birefringence-like 38 | May act as a bridging protein that binds pectin and other cell wall polysaccharides. Probably involved in maintaining | | drought stress response | Shuai, P., Liang, D., Zhang, Z., Yin, W., & Xia, X. (2013). Identification of drought-responsive and novel Populus trichocarpamicroRNAs by high-throughput sequencing and their |

| | | | | | | | | | |
|---|---|---|---|---|---|---|---|---|---|
| | | | | | | esterification of pectins | | | targets using degradome analysis. Bmc Genomics, 14(1), 233. |
| 8.g199 | KAF3962723 | hypothetical protein CMV_012798 | *Castanea mollissima* | | | | | | |
| 8.g2624 | No hits found | | | | | | | | |
| 8.g2764 | RUS78569 | hypothetical protein EGW08_013676, partial | *Elysia chlorotica* | | | | | | |
| 8.g2926 | RUS32681 | hypothetical protein BC938DRAFT_474617 | *Jimgerdemannia flammicorona* | | | | | | |
| 8.g3326 | XP_030953364 | sister chromatid cohesion protein SCC2 | *Quercus lobata* | SCC2_ARATH | sister chromatid cohesion protein SCC2 | Essential protein required for cell fate determination during embryogenesis | Sebastian, J., Ravi, M., Andreuzza, S., Panoli, A. P., Marimuthu, M. P., & Siddiqi, I. (2009). The plant adherin AtSCC2 is required for embryogenesis and sister-chromatid cohesion during meiosis in Arabidopsis. The Plant Journal, 59(1), 1-13. | | |

| | | | | | | | | | |
|---|---|---|---|---|---|---|---|---|---|
| 8.g3872 | XP_030953755 | surfeit locus protein 2 | *Quercus lobata* | SURF1_ARATH | Surfeit locus protein 1 | Probably involved in the biogenesis of the COX complex | | | |
| 8.g4454 | KAF3955069 | hypothetical protein CMV_019673 | *Castanea mollissima* | | | | | | |
| 8.g4554 | KAF3975054 | hypothetical protein CMV_001659 | *Castanea mollissima* | | | | | | |
| 8.g935 | KAF3972759 | hypothetical protein CMV_003766 | *Castanea mollissima* | | | | | | |
| 9.g1013 | KAF3963767 | hypothetical protein CMV_011877 | *Castanea mollissima* | | | | | | |
| 9.g1025 | XP_023873320 | cytosolic sulfotransferase 5-like | *Quercus suber* | | | | | | |
| 9.g2127 | No hits found | | | | | | | | |
| 9.g4398 | KAB5561320 | hypothetical protein DKX38_006277 | *Salix brachista* | | | | | | |
| 9.g4504 | KAF3976808 | hypothetical protein CMV_000062 | *Castanea mollissima* | TBL33_ARATH | Protein trichome birefringence-like 33 | Probable xylan acetyltransferase that plays a role in xylan acetylation and normal deposition of secondary cell walls | Yuan, Y., Teng, Q., Zhong, R., Haghighat, M., Richardson, E. A., & Ye, Z. H. (2016). Mutations of Arabidopsis TBL32 and TBL33 affect xylan acetylation and secondary | drought stress response | Shuai, P., Liang, D., Zhang, Z., Yin, W., & Xia, X. (2013). Identification of drought-responsive and novel Populus trichocarpamicroRNAs by high-throughput sequencing and their targets using degradome analysis. Bmc Genomics, 14(1), |

| | | | | | | | | | |
|---|---|---|---|---|---|---|---|---|---|
| | | | | | | | wall deposition. PLoS One, 11(1), e0146460. | | 233. |
| 9.g4548 | XP_023928861 | uncharacterized protein LOC112040195 | *Quercus suber* | | | | | | |
| 9.g4606 | XP_030928366 | LOW QUALITY PROTEIN: E3 ubiquitin-protein ligase SHPRH | *Quercus lobata* | | | | | | |
| 9.g653 | KAF3952878 | hypothetical protein CMV_021616 | *Castanea mollissima* | | | | | | |
| 10.g4171 | ONI05229 | hypothetical protein PRUPE_6G363600 | *Prunus persica* | | | | | | |
| 11.g2603 | XP_023892377 | amino acid permease 6-like | *Quercus suber* | AAP6_ARATH | Amino acid permease 6 | Amino acid-proton symporter. Stereospecific transporter with a broad specificity for tryptophan, proline, and neutral and acidic amino acids. | Rentsch, D., Hirner, B., Schmelzer, E., & Frommer, W. B. (1996). Salt stress-induced proline transporters and salt stress-repressed broad specificity amino acid permeases identified by suppression of | environmental stress response | Rentsch, D., Hirner, B., Schmelzer, E., & Frommer, W. B. (1996). Salt stress-induced proline transporters and salt stress-repressed broad specificity amino acid permeases identified by suppression of a yeast amino acid permease-targeting mutant. The Plant Cell, 8(8), 1437-1446. |

| | | | | | | | | | |
|---|---|---|---|---|---|---|---|---|---|
| | | | | | | | a yeast amino acid permease-targeting mutant. The Plant Cell, 8(8), 1437-1446. | | |
| 11.g787 | No hits found | | | | | | | | |
| 12.g1692 | KAF3972258 | hypothetical protein CMV_004214 | *Castanea mollissima* | | | | | | |
| 12.g2592 | No hits found | | | | | | | | |
| 12.g3049 | XP_023886583 | transcription factor TCP15 | *Quercus suber* | TCP15_ARATH | transcription factor TCP15 | Transcription factor involved the regulation of plant development. Together with TCP14, modulates plant stature by promoting cell division in young internodes. | Kieffer, M., Master, V., Waites, R., & Davies, B. (2011). TCP14 and TCP15 affect internode length and leaf shape in Arabidopsis. The Plant Journal, 68(1), 147-158. | environmental stress response | Viola, I. L., Camoirano, A., & Gonzalez, D. H. (2016). Redox-dependent modulation of anthocyanin biosynthesis by the TCP transcription factor TCP15 during exposure to high light intensity conditions in Arabidopsis. Plant Physiology, 170(1), 74-85. |
| 12.g922 | KAF3957893 | hypothetical protein CMV_017140 | *Castanea mollissima* | | | | | | |

Suppl. Table 5. List of 20 most informative SNPs as selected by the eSPA Analysis allowing for 85% correct classification.

| | scaffold | position | gene | swiss_prot_id_(arabidopsis_thaliana) |
|---|---|---|---|---|
| 41 | Fsyl.bwa_aln.clean.counts_GATC.12g6 | 1385435 | / | / |
| 56 | Fsyl.bwa_aln.clean.counts_GATC.12g7 | 31456694 | Fsyl.bwa_aln.clean.counts_GATC.12g7.g3617.t1 | Q9S7R3 |
| 14 | Fsyl.bwa_aln.clean.counts_GATC.12g12 | 7252002 | / | / |
| 9 | Fsyl.bwa_aln.clean.counts_GATC.12g11 | 21712586 | / | / |
| 42 | Fsyl.bwa_aln.clean.counts_GATC.12g6 | 1559115 | / | / |
| 43 | Fsyl.bwa_aln.clean.counts_GATC.12g6 | 1559277 | / | / |
| 44 | Fsyl.bwa_aln.clean.counts_GATC.12g6 | 1559370 | / | / |
| 27 | Fsyl.bwa_aln.clean.counts_GATC.12g2 | 22433752 | / | / |
| 58 | Fsyl.bwa_aln.clean.counts_GATC.12g7 | 33110000 | Fsyl.bwa_aln.clean.counts_GATC.12g7.g3816.t1 | Q94ID6 |
| 36 | Fsyl.bwa_aln.clean.counts_GATC.12g4 | 34077017 | Fsyl.bwa_aln.clean.counts_GATC.12g4.g3980.t1 | Q67YU0 |
| 84 | Fsyl.bwa_aln.clean.counts_GATC.12g9 | 38251152 | / | / |
| 85 | Fsyl.bwa_aln.clean.counts_GATC.12g9 | 38652380 | / | / |
| 59 | Fsyl.bwa_aln.clean.counts_GATC.12g8 | 1582675 | / | / |
| 1 | Fsyl.bwa_aln.clean.counts_GATC.12g1 | 10993370 | / | / |
| 78 | Fsyl.bwa_aln.clean.counts_GATC.12g9 | 20434572 | / | / |
| 82 | Fsyl.bwa_aln.clean.counts_GATC.12g9 | 37955715 | Fsyl.bwa_aln.clean.counts_GATC.12g9.g4504.t1 | F4IH21 |
| 80 | Fsyl.bwa_aln.clean.counts_GATC.12g9 | 25538827 | Fsyl.bwa_aln.clean.counts_GATC.12g9.g3080.t1 | Q9FMK9 |
| 34 | Fsyl.bwa_aln.clean.counts_GATC.12g4 | 33216309 | / | / |
| 70 | Fsyl.bwa_aln.clean.counts_GATC.12g9 | 5021966 | / | / |
| 69 | Fsyl.bwa_aln.clean.counts_GATC.12g8 | 38060740 | / | / |

**Suppl. Info 1**. Software pipeline and commands used for PoolSeq analysis.

```
#AUTOTRIM
perl ../autotrim-master/autotrim.pl -fofn files.txt -trim ../trimmomatic_options -tt 12 -log /data/FagusPools -tp /home/mpfenninger/Trimmomatic-0.39/trimmomatic-0.39.jar -fqcp /home/mpfenninger/FastQC/fastqc

#Mapping with BWA

bwa index Beech_12Chr.masked.fas

bwa mem -t 12 -k 30 /Genome/Fagus_sylvatica_genome.fasta XXX_R1.autotrim.paired.fq XXX_R2_autotrim.paired.fq > XXX_bwamem.sam

while read poollist
do samtools view -b $poollist"_bwamem.sam" > $poollist".bam"
done < poollist
wait

samtools sort XX.bam > XX.sort.bam

#Remove Duplicates with Picard

java -jar /home/mpfenninger/Picard/picard.jar MarkDuplicates I=MPG.sort.bam O=MPG.sort.rmd.bam M=PoolMPG.dupstat.txt VALIDATION_STRINGENCY=SILENT REMOVE_DUPLICATES=TRUE

#Remove low quality mappings with SAMtools

samtools view -q 20 -@ 14 -f 0x0002 -F 0x0004 -F 0x0008 -b MPG.sort.rmd.bam > MPG.sort.rmd.q20.bam

samtools index MPG.sort.rmd.q20.bam

#Create pileup with mpileup

samtools mpileup -f /data/FagusPools/Genome/Fagus_sylvatica_genome.fasta -B -Q 0 MPG.sort.rmd.q20.bam MPK.sort.rmd.q20.bam
```

```
SPG.sort.rmd.q20.bam SPK.sort.rmd.q20.bam > FagusPool.mpileup

#Convert mpileup to sync with Popoolation 2_1201

java -jar ~/popoolation2_1201/mpileup2sync.jar --input FagusPool.mpileup --output FagusPool.sync --fastq-type sanger --min-qual 20 --threads 14

#Filtering for indels
#get indels
perl ~/popoolation2_1201/indel_filtering/identify-indel-regions.pl --indel-window 5 --input FagusPool.mpileup --output FagusPool.indels.gtf

#remove indels from sync
perl ~/popoolation2_1201/indel_filtering/filter-sync-by-gtf.pl --input FagusPool.sync --gtf FagusPool.indels.gtf --output FagusPool.idf.sync


#Fst calculation

perl ~/popoolation2_1201/fst-sliding.pl --input FagusPool.idf.sync --output Fagus.fst --min-count 2 --min-coverage 15 --max-coverage 2% --pool-size 100 --window-size 1000 --step-size 1000

#Fisher's exact test with PoolSeq 0.35 in R
library("poolSeq")
Fagus <- read.sync(file="/data/FagusPools/FagusPool.idf.sync", gen=c(0,1,0,1), repl=c(1,1,2,2),  keepOnlyBiallelic=TRUE)
afMPG <- af(sync=Fagus, repl=1, gen=0)
afMPK <- af(sync=Fagus, repl=1, gen=1)
afSPG <- af(sync=Fagus, repl=2, gen=0)
afSPK <- af(sync=Fagus, repl=2, gen=1)

write.table(afMPG, "afMPG.txt", sep="\t")
write.table(afMPK, "afMPK.txt", sep="\t")
write.table(afSPG, "afSPG.txt", sep="\t")
write.table(afSPK, "afSPK.txt", sep="\t")

covMPG <- coverage(sync=Fagus, repl=1, gen=0)
```

```
covMPK <- coverage(sync=Fagus, repl=1, gen=1)
covSPG <- coverage(sync=Fagus, repl=2, gen=0)
covSPK <- coverage(sync=Fagus, repl=2, gen=1)

AFC_MP <- afMPG - afMPK
write.table(AFC_MP, "AFC_MP.txt", sep="\t")
AFC_SP <- afSPG - afSPK
write.table(AFC_SP, "AFC_SP.txt", sep="\t")

NA_MPG <- t(covMPG * afMPG)
Na_MPG <- covMPG - NA_MPG
NA_MPK <- t(covMPK * afMPK)
Na_MPK <- covMPK - NA_MPK

NA_SPG <- t(covSPG * afSPG)
Na_SPG <- covSPG - NA_SPG
NA_SPK <- t(covSPK * afSPK)
Na_SPK <- covSPK - NA_SPK

p.valuesMPmaxcov120 <- chi.sq.test(A0 = NA_MPG, a0 = Na_MPG, At = NA_MPK, at = Na_MPK, min.cov=15, min.cnt = 3, max.cov=120, log =
TRUE)
quantile(p.valuesMPmaxcov120, probs = c(0.999, 0.9999, 0.99999), na.rm = TRUE, names = TRUE)
p.valuesSPmaxcov120 <- chi.sq.test(A0 = NA_SPG, a0 = Na_SPG, At = NA_SPK, at = Na_SPK, min.cov=15, min.cnt = 3, max.cov=120, log =
TRUE)
quantile(p.valuesMPmaxcov120, probs = c(0.999, 0.9999, 0.99999), na.rm = TRUE, names = TRUE)

pdf("MPmaxcov120.pdf")
plot(p.valuesMPmaxcov120, main=paste0("MP"), ylim=c(0, max(p.valuesMPmaxcov120, na.rm=TRUE)), xlab="position", ylab="-log10(p)", pch=".")
dev.off()
pdf("SPmaxcov120.pdf")
plot(p.valuesSPmaxcov120, main=paste0("SP"), ylim=c(0, max(p.valuesMPmaxcov120, na.rm=TRUE)), xlab="position", ylab="-log10(p)", pch=".")
dev.off()
```

```
write.table(p.valuesMPmaxcov120, "p.valuestotmaxcov120.txt", sep="\t")
write.table(p.valuestotmaxcov120, "p.valuestotmaxcov120.txt", sep="\t")

#cmh-Test

covG <- t(coverage(sync=Fagus, repl=1:2, gen=0))
covK <- t(coverage(sync=Fagus, repl=1:2, gen=1))
A_G <- t(af(sync=Fagus, repl=1:2, gen=0)) * covG
a_G <- covG - A_G
write.table(A_G, "afG.txt", sep="/t")
A_K <- t(af(sync=Fagus, repl=1:2, gen=1)) * covK
a_K <- covK - A_K
write.table(A_K, "afK.txt", sep="/t")
p.valuestot <- cmh.test(A0 = A_G, a0=a_G, At=A_K, at=a_K, min.cov=15, max.cov=100, min.cnt=3, log=TRUE)
write.table(p.valuestot, "p.valuestot.txt", sep="\t")




p.valuestot <- cmh.test(A0 = A_G, a0=a_G, At=A_K, at=a_K, min.cov=15, max.cov=100, min.cnt=3, log=TRUE)
pdf("cmh.pdf")
plot(p.valuestot, main=paste0("cmh_fdr"), ylim=c(0, max(p.valuestot, na.rm=TRUE)), xlab="position", ylab="-log10(p)", pch=".")
dev.off()
quantile(p.valuestot, probs = c(0.999, 0.9999, 0.99999), na.rm = TRUE, names = TRUE)
write.table(p.valuestot, "p.valuestot.txt", sep="\t")
write.table(A_G, "afG.txt", sep="/t")
write.table(A_K, "afK.txt", sep="/t")
length(p.fdr[p.fdr > 4])
```

**Suppl. Info 2**. Workflow individual reseq GWAS
####workflow beech GWAS individual reseq data######
####May 2020 Barbara Feldmeyer#######
autotrim v.0.6.1
bcftools v.1.9
gatk v.4.1.7.0
picard v.2.20.8
plink v.1.90b6.13
samtools v.1.10

1) trim reads with autotrim (https://github.com/schellt/autotrim)

2) index genome file

    bwa index Fagus_sylvatica_genome_v2_masked.fasta
    samtools faidx Fagus_sylvatica_genome_v2_masked.fasta

AND create dictionary

    picard CreateSequenceDictionary.jar R=Fagus_sylvatica_genome_v2_masked.fasta

Step3a map reads to genome Fagus_sylvatica_genome_v2_masked.fasta

    bwa mem -M -t 10 Fagus_sylvatica_genome_v2_masked.fasta $i_1_autotrim.paired.fq $i_2_autotrim.paired.fq | samtools sort -l 9 -O BAM -o ${FILES[$SLURM_ARRAY_TASK_ID]}.sort.bam"#

3b)MarkDuplicates

    picard MarkDuplicates I=$i.sort.bam O=$i.sort.bam_marked_dup.bam M=$i.sort.bam_marked_dup_metrics.txt VALIDATION_STRINGENCY=SILENT REMOVE_DUPLICATES=true

3c) SortSAM

    picard SortSam I=$i.sort.bam_marked_dup.bam O=$i.sort.bam_marked_dup.bam_sorted.bam VALIDATION_STRINGENCY=SILENT SORT_ORDER=coordinate

3d) index bam

```
picard BuildBamIndex INPUT=$i.sort.bam_marked_duplicates.bam_sorted.bam
```

3e) check mapping quality with qualimap

```
qualimap multi-bamqc -r -d qualimap_commands
```

==>> PF_001 very bad mapping quality (<1%!!!!) and skewed GC ratio in PF_050 => remove these two samples from further analyses

3f) create file with sample.bam specific read groups

```
sed 's/.*data\///' checkSize.xls | sed 's/\//\tRGSM=/' |sed 's/_BD.*HW/ RGID=HW/'| sed 's/_L/./'| sed 's/_[0-9].fq.gz/ RGLB=SOME RGPL=illumina RGPU=2/' > readgroups_beech.txt
```

3g) create batch file to run picard to modify and add readgroups to .bam file header

```
picard AddOrReplaceReadGroups I="$i.sort.bam_marked_duplicates.bam_sorted.bam" O="$i.sort.bam_marked_duplicates_sorted_rehead.bam" RGSM=$i RGID=HWTH2DSXX.4 RGLB=SOME RGPL=illumina RGPU=2" &
```

3h) index renamed bams

```
##########
#Round1
##########
```

4a)run HaplotypeCaller in GVCF mode

```
gatk HaplotypeCaller -I $i -O $i_haploCall.gvcf -R Fagus_sylvatica_genome.fasta -ERC GVCF
```

4b) combine using combineGVCF

```
gatk CombineGVCFs -R Fagus_sylvatica_genome_v2_masked.fasta --variant PF_002.sort.bam_marked_duplicates_sorted_rehead.bam_haploCall.gvcf --variant PF_003.bam_marked_duplicates_sorted_rehead.bam_haploCall.gvcf ... -O beech_cohort98indivs_R1.gvcf" &
```

4c) GenotypeGVCFs to jointly call Haplotypes

```
GenotypeGVCFs -R  Fagus_sylvatica_genome_v2_masked.fasta -V beech_cohort98indivs_R1.gvcf -O beech_cohort98indivs_R1_genotyped.gvcf
```

4d) call and subset SNPs

gatk SelectVariants -V beech_cohort98indivs_R1_genotyped.gvcf --select-type-to-include SNP -O beech_cohort98indivs_R1_genotyped_SNPs.vcf

4e) create summary stats
bcftools stats beech_cohort98indivs_R1_genotyped_SNPs.vcf > beech_cohort98indivs_R1_genotyped_SNPs_summaryStats.txt

4f) hard-filter SNPs (we conduct this conservative hard filtration step since we don't have any pre-existing SNP set available to recalibrate SNPs)
gatk VariantFiltration -R Fagus_sylvatica_genome_v2_masked.fasta -V beech_cohort98indivs_R1_genotyped_SNPs.vcf -O beech_cohort98indivs_R1_genotyped_SNPs_hardfiltration.vcf --filter-expression 'QD < 2.0' --filter-name 'QD2' --filter-expression 'MQ < 50.0' --filter-name 'MQ50' --filter-expression 'MQRankSum < -12.5' --filter-name 'MQRankSum-12.5' --filter-expression 'ReadPosRankSum < -8.0' --filter-name 'ReadPosRankSum-8' --filter-expression 'FS > 80.0' --filter-name 'FS' --filter-expression 'SOR > 4.00' --filter-name 'SOR_4' --filter-expression 'QUAL < 10.0' --filter-name 'QUAL_10'

-grep and save passed variants
grep -E '^#|PASS' beech_ cohort98indivs_R1_genotyped_SNPs_hardfiltration.vcf > beech_cohort98indivs_R1_genotyped_SNPs_hardfiltrationPASS.vcf

4g) Baserecalibration
gatk --java-options "-Xmx4g" BaseRecalibrator -I $i -O fagus_recalR1_data.table -R Fagus_sylvatica_genome_v2_masked.fasta --known-sites beech_cohort98indivs_R1_genotyped_indels_hardfiltration.vcf --known-sites beech_cohort98indivs_R1_genotyped_SNPs_hardfiltration.vcf

4h) apply BQSR
gatk --java-options "-Xmx4g" ApplyBQSR -I $i -O $i_abqsr_R1.bam -R Fagus_sylvatica_genome_v2_masked.fasta --bqsr-recal-file fagus_recalR1_data.table

###########################
#Round 2
##############################
5a) R2_haplocaller
gatk --java-options "-Xmx4g" HaplotypeCaller -I $i -O $i_haploCall_R2.gvcf -R Fagus_sylvatica_genome_v2_masked.fasta -ERC GVCF

5b) R2_combine GVCF
gatk CombineGVCFs -R Fagus_sylvatica_genome_v2_masked.fasta --variant .... -O beech_cohort98indivs_R2.gvcf" &

5c) R2_GenotypeGVCFs

```
gatk --java-options "-Xmx250G" GenotypeGVCFs -R Fagus_sylvatica_genome_v2_masked.fasta -V beech_cohort98indivs_R2.gvcf -O beech_cohort98indivs_R2_genotyped.gvcf
```

5d) R2_call, select and subset SNPs

```
gatk SelectVariants -V beech_cohort98indivs_R2_genotyped.gvcf --select-type-to-include SNP -O beech_cohort98indivs_R2_genotyped_SNPs.vcf
```

create summary stats

```
bcftools stats beech_cohort98indivs_R2_genotyped_SNPs.vcf > beech_cohort98indivs_R2_genotyped_SNPs_summaryStats.txt
```

6) variant filtration

6a) Variants to table (Extract Variant Quality Score)

```
gatk --java-options "-Xmx250G" VariantsToTable R Fagus_sylvatica_genome_v2_masked.fasta -V beech_cohort98indivs_R2_genotyped_SNPs.vcf -F CHROM -F POS -F QUAL -F QD -F DP -F MQ -F MQRankSum -F FS -F ReadPosRankSum -F SOR -O cohort_all98beechSamples_genotyped_snp.table" &
```

6b)create diagnostic plots for Variants run skript in R following (https://evodify.com/gatk-in-non-model-organism/)

```
########################################
library('gridExtra')
library('ggplot2')

VCFsnps <- read.csv('cohort_all98beechSamples_genotyped_snp.table', header = T, na.strings=c("","NA"), sep = "\t")
VCFindel <- read.csv('cohort_all98beechSamples_genotyped_indel.table', header = T, na.strings=c("","NA"), sep = "\t")
dim(VCFsnps)
dim(VCFindel)
VCF <- rbind(VCFsnps, VCFindel)
VCF$Variant <- factor(c(rep("SNPs", dim(VCFsnps)[1]), rep("Indels", dim(VCFindel)[1])))

snps <- '#A9E2E4'
indels <- '#F4CCCA'

DP <- ggplot(VCF, aes(x=DP, fill=Variant)) + geom_density(alpha=0.3) +
  geom_vline(xintercept=c(10,6200))

QD <- ggplot(VCF, aes(x=QD, fill=Variant)) + geom_density(alpha=.3) +
```

```
  geom_vline(xintercept=2, size=0.7)

FS <- ggplot(VCF, aes(x=FS, fill=Variant)) + geom_density(alpha=.3) +
  geom_vline(xintercept=c(60, 200), size=0.7) + ylim(0,0.1)

MQ <- ggplot(VCF, aes(x=MQ, fill=Variant)) + geom_density(alpha=.3) +
  geom_vline(xintercept=40, size=0.7)

MQRankSum <- ggplot(VCF, aes(x=MQRankSum, fill=Variant)) + geom_density(alpha=.3) +
  geom_vline(xintercept=-20, size=0.7)

SOR <- ggplot(VCF, aes(x=SOR, fill=Variant)) + geom_density(alpha=.3) +
  geom_vline(xintercept=c(4, 10), size=1, colour = c(snps,indels))

ReadPosRankSum <- ggplot(VCF, aes(x=ReadPosRankSum, fill=Variant)) + geom_density(alpha=.3) +
  geom_vline(xintercept=c(-10,10,-20,20), size=1, colour = c(snps,snps,indels,indels)) + xlim(-30, 30)

svg("cohort_98individuals_R2_beech20200609.svg", height=20, width=15)
theme_set(theme_gray(base_size = 18))
grid.arrange(QD, DP, FS, MQ, MQRankSum, SOR, ReadPosRankSum, nrow=4)
dev.off()
###########################
```

6c) VariantFiltration

```
gatk VariantFiltration -R Fagus_sylvatica_genome_v2_masked.fasta -V beech_cohort98indivs_R2_genotyped_SNPs.vcf -O beech_cohort98indivs_R2_genotyped_SNPs.vcf_filtered.vcf --filter-expression 'QD < 2.0' --filter-name 'QD2' --filter-expression 'MQ < 40.0' --filter-name 'MQ_40' --filter-expression 'SOR > 4.00' --filter-name 'SOR_4' --filter-expression 'QUAL < 10.0' --filter-name 'QUAL_10' --filter-expression 'FS > 60.0' --filter-name 'FS_60'
```

-grep and save passed variants

```
grep -E '^#|PASS' beech_cohort98indivs_R2_genotyped_SNPs.vcf_filtered.vcf > beech_cohort98indivs_R2_genotyped_SNPs.vcf_filteredPASSED.vcf
```

-stats

```
bcftools stats beech_cohort98indivs_R2_genotyped_SNPs.vcf_filteredPASSED.vcf
```

7) Plink association analysis

7.1. create phenotype file containing relevant phenoype information

beech_phenotype.phe

7.2. convert .vcf file to plink .ped format and keep biallic only

```
plink --vcf beech_cohort98indivs_R2_genotyped_SNPs.vcf_filteredPASSED_genotypeFiltered.vcf --biallelic-only strict --double-id --allow-extra-chr --set-missing-var-ids @:# --indep-pairwise 50 10 0.1 --out beech_cohort98indivs_SNPs
```

7.2b. additionally create a file with all sites, i.e. do not remove linked loci

```
plink --vcf beech_cohort98indivs_R2_genotyped_SNPs.vcf_filteredPASSED_genotypeFiltered.vcf --biallelic-only strict --double-id --allow-extra-chr --set-missing-var-ids @:# --make-bed --out beech_cohort98indivs_linkedSNPs
```

7.3. create PCA from linkage-pruned sites

-run plink to create PCA values

```
plink --vcf beech_cohort98indivs_R2_genotyped_SNPs.vcf_filteredPASSED_genotypeFiltered.vcf --double-id --allow-extra-chr --set-missing-var-ids @:# --extract beech_cohort98indivs_SNPs.prune.in --make-bed --pca --out beech_cohort98indivs_SNPs
```

-run R to plot the PCA

```
library(tidyverse)
pca<-read_table2("beech_cohort98indivs_SNPs.eigenvec", col_names=FALSE)
eigenval<-scan("beech_cohort98indivs_SNPs.eigenval")
 pca <- pca[,-1]
 names(pca)[1] <- "ind"
 names(pca)[2:ncol(pca)] <- paste0("PC", 1:(ncol(pca)-1))
```

7.4. create missing stats

```
plink -bfile beech_cohort98indivs_SNPs --missing --out miss_stat_SNPs --allow-extra-chr
```

7.5. summary stats allele frequency

```
plink -bfile beech_cohort98indivs_SNPs --freq --out freq_stat_SNPs --allow-extra-chr
```

7.6. association analysis with automatic correction for multiple testing

```
plink --assoc --bfile beech_cohort98indivs_SNPs --allow-no-sex --adjust --allow-extra-chr
```
